# Pathologic α-Synuclein Species Activate LRRK2 in Pro-Inflammatory Monocyte and Macrophage Responses

**DOI:** 10.1101/2020.05.04.077065

**Authors:** Enquan Xu, Ravindra Boddu, Hisham A. Abdelmotilib, Kaela Kelly, Arpine Sokratian, Ashley S. Harms, Aubrey M. Schonhoff, Nicole Bryant, Irene E. Harmsen, Michael G. Schlossmacher, Sidhanth Chandra, Valentina Krendelshchikova, Zhiyong Liu, Andrew B. West

## Abstract

Missense mutations in the *LRRK2* gene that lead to LRRK2 kinase hyperactivity can cause Parkinson’s disease (PD). The link between LRRK2 and α-synuclein aggregation in PD remains enigmatic. Numerous reports suggest critical LRRK2 functions in microglial responses. Herein, we find that LRRK2-positive immune cells in the brain represent CD68-positive pro-inflammatory, monocyte-derived macrophages, distinct from microglia. Rod α-synuclein fibrils stimulate LRRK2 kinase activity in monocyte-derived macrophages, and LRRK2 mutations lead to enhanced recruitment of classical monocytes into the midbrain in response to α-synuclein. LRRK2 kinase inhibition blocks α-synuclein fibril induction of LRRK2 protein in both human and murine macrophages, with human cells demonstrating much higher LRRK2 levels and kinase activity than equivalent murine cells. Further, interferon-γ strongly induces LRRK2 kinase activity in primary human macrophages in comparison to weak effects observed in murine cells. These results highlight peripheral immune responses in LRRK2-linked paradigms that further connect two central proteins in PD.

## Introduction

Missense mutations in autosomal-dominant Parkinson’s disease (PD) in the *leucine-rich repeat kinase 2* (*LRRK2*) gene are one of the most common known genetic causes of PD and neurodegeneration (Trinh et al., 2014; West, 2017). Pathogenic *LRRK2* mutations lead to upregulated LRRK2 autophosphorylation nearby the LRRK2 ROC Rab-like domain at Ser1292 (Sheng et al., 2012) as well as upregulated phosphorylation of Rab substrates, notably Rab10 (Steger et al., 2016). *LRRK2* is independently associated with PD through genome-wide association studies, in addition to associations with irritable bowel syndrome and mycobacterium infection susceptibility (Fava et al., 2016; Umeno et al., 2011; Zhang et al., 2009). Numerous studies in different models suggest that LRRK2 expression in neurons affects intrinsic vulnerability to α-synuclein aggregation (Bae et al., 2018; Bieri et al., 2019; Lin et al., 2009; Volpicelli-Daley et al., 2016; Zhao et al., 2017). We previously detailed LRRK2 protein expression in healthy mouse, rat, and human brains, identifying distinct neuronal subtypes with high LRRK2 protein levels (Davies et al., 2013; West et al., 2014).

The distribution of LRRK2-positive cells in the mammalian brain changes under conditions of neuroinflammation. In rats, *Lrrk2* expression occurs in some CD68+ cells that are not present in healthy brains but induced by rAAV2-α-synuclein expression or the toll-like receptor (TLR)-4 agonist lipopolysaccharide (LPS, (Daher et al., 2014)). Aggregated α-synuclein protein serves as a strong pro-inflammatory agonist in different types of myeloid cells, including monocytes and microglia, possibly through interactions with TLRs (Fellner et al., 2013; Grozdanov et al., 2019; Kim et al., 2018a; Kwon et al., 2019). In cultured mouse macrophages, Lrrk2 deficiency and its kinase inhibition specifically impair chemotaxis without apparent effect on secreted cytokines and chemokines (Levy et al., 2020; Moehle et al., 2015; Moehle et al., 2012; Shutinoski et al., 2019). Recent studies have demonstrated that *Lrrk2* knockout or kinase inhibition diminishes innate immune cell responses in the brain, as evidenced by weakened microglial responses to LPS (Daher et al., 2014; Dwyer et al., 2020; Kim et al., 2012; Ma et al., 2016; Moehle et al., 2012; Russo et al., 2019), to transgenic mutant A53T-α-synuclein expression (Lin et al., 2009), to rAAV2-α-synuclein expression (Daher et al., 2014), to HIV-1 TAT peptide injections (Puccini et al., 2015), and in the context of experimental uveitis (Wandu et al., 2015). Based on these results, LRRK2 has been proposed as a key mediator of innate immune responses in the brain in disease (Russo et al., 2014; Schapansky et al., 2015).

In rodent models of PD, peripheral immune cells interact with resident immune cells in diseased tissue (Tansey and Romero-Ramos, 2019). In the brain, resident macrophages of the innate immune system are stable populations of cells that arise from yolk sac precursors during embryonic development (Ginhoux et al., 2010). Resident brain immune cells are composed in-part of parenchymal microglia as well as non-parenchymal macrophages in the perivasculature and meningeal spaces (Goldmann et al., 2016). In disease, bone marrow-derived monocytes can be recruited to the brain and differentiate into macrophages, although there is less known about this process, in part because there are no effective markers that might reliably delineate monocyte-derived macrophages from other immune cells in the context of disease. While it might be presumed that the intrinsic action of LRRK2 in resident microglia in the brain have driven immunological phenotypes observed in numerous past studies, surprisingly, there have been no previous concerted efforts to specifically localize LRRK2 protein expression and kinase activity to microglia cells in the brain, or to distinguish their action from other types of interacting immune cells. We postulate that the lack of knowledge of LRRK2 action within the immune system may hinder the development of therapeutics targeting LRRK2 for neuroprotection. Herein, we find that pathological α-synuclein fibrils activate LRRK2 in immune cells, and that this process is conserved from mouse to human; moreover, we discovered that LRRK2 protein levels, kinase activity, and α-synuclein induction may be confined to monocyte-derived macrophages that are distinct from microglia and their activated progeny. Moreover, our results highlight species differences between murine and human cells regarding their degree of LRRK2 activity in α-synuclein stimulation as well as responsiveness to interferon (IFN)γ signaling pathways.

## Results

### α-Synuclein fibrils activate LRRK2 in mouse and human monocyte-derived macrophages

Recent data from primary human monocytes and mouse microglia cell line cultures (BV2 from C57BL/6 mice) have shown that rod-shaped α-synuclein fibrils induce pro-inflammatory responses, in contrast to α-synuclein oligomers, ribbons thereof, or monomeric protein species, that all lacked these effects (Grozdanov et al., 2019). To determine whether rod α-synuclein fibrils might affect LRRK2 expression and kinase activity towards the LRRK2 kinase substrate Rab10 that is highly expressed in immune cells (Liu et al., 2020), we first created a panel of carefully curated mouse or human α-synuclein recombinant proteins, with monomer preparations that matched spontaneously fibrilized rod-type fibrils, sonicated to increase solubility to an approximate homogeneity in length (20 nm, Figure 1A-C). In CSF1-expanded (macrophage colony-stimulating factor, M-CSF) mouse bone marrow-derived primary macrophages, application of ∼642 pM in-solution of mouse α-synuclein rod fibril particles (or 1 μg per mL of ∼20 nM rod-fibrils) led to an induction of Lrrk2 protein expression (∼2.5 fold, Figure 1D,E) as well as phosphorylation of the Lrrk2 kinase substrate Rab10 (∼2 fold, Figure 1D,F). Exposure to 1 μg per mL of monomeric α-synuclein (∼69 nM) lacked this effect. Notably, inclusion of a relatively selective LRRK2 kinase inhibitor, MLi2, at low nanomolar concentrations (100 nM), completely blocked both Rab10 phosphorylation as well as the fibril-induction of total Lrrk2 protein. In macrophages without α-synuclein fibril stimulation, MLi2 exposure did not reduce levels of total Lrrk2 protein in controls, or after incubation with monomers, suggesting that the inhibitor itself does not reduce Lrrk2 levels. MLi2, as opposed to older generations of kinase inhibitors used in past studies of immune cells (Moehle et al., 2015), is known not to bind to or inhibit other known kinases or proteins at concentrations of less than 100 nM, with equal potency in blocking mouse Lrrk2 and human LRRK2 kinase activities (Fell et al., 2015; Kelly et al., 2018; Volpicelli-Daley et al., 2016).

**Figure 1.**
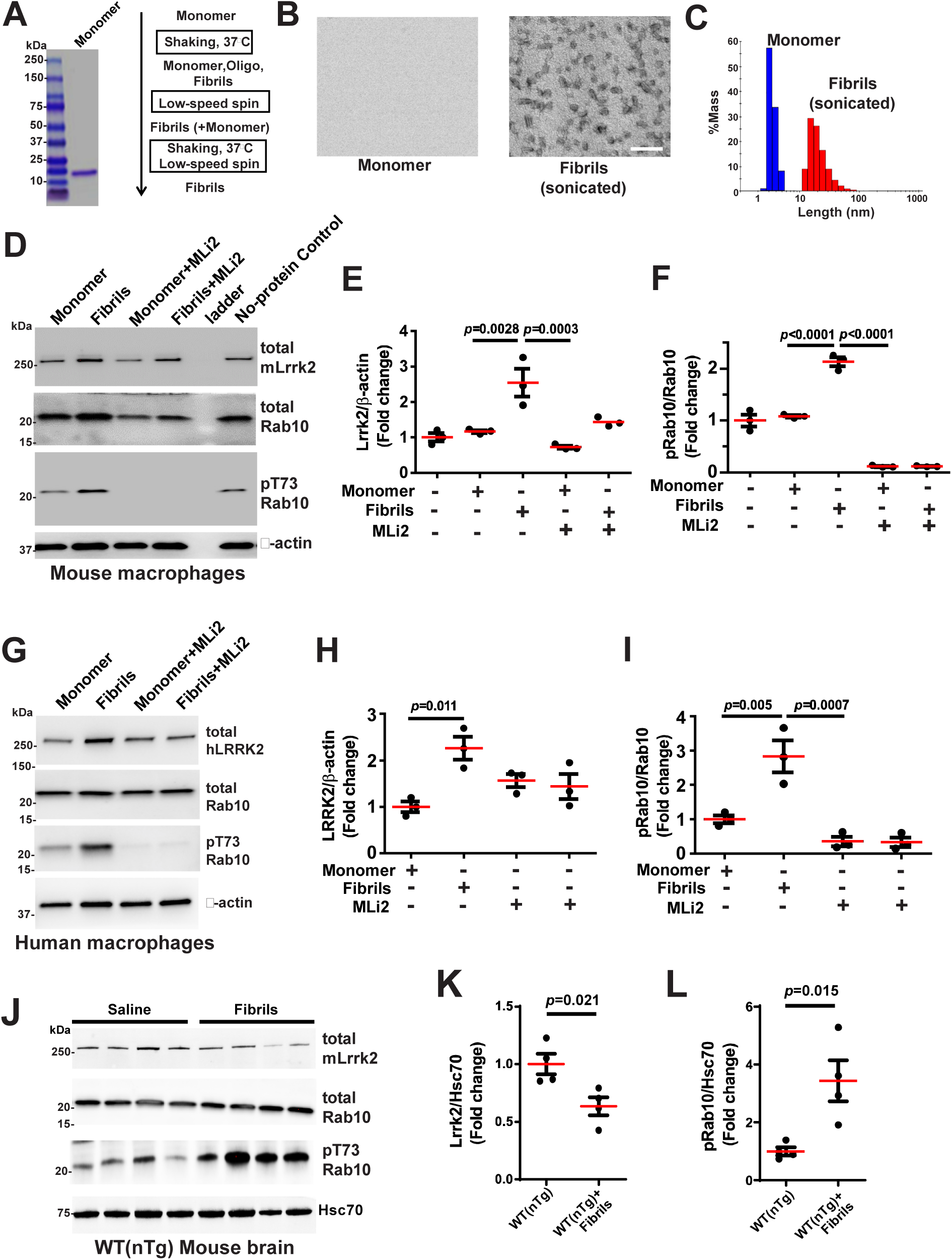
α-Synuclein fibrils induce LRRK2 activity in monocyte-derived macrophages. **a.** α-Synuclein purification and summary of fibril isolation. **b.** Electron-microscopy photomicrographs of monomer and sonicated fibrils. Scale bar is 100 nm. **c.** Light-scattering characterization of matched monomer and fibril preparations. **d.** Representative immunoblots of mouse macrophages, and **e., f.**, their quantification, from lysates of primary mouse bone marrow-derived macrophage cells from adult male WT(nTg, non-transgenic, C57BL/6J) mice, polarized with mouse M-CSF (macrophage-colony stimulating factor). Macrophages were treated with 1 μg mL^-1^ of mouse monomer or fibril protein, with or without pretreatment of the LRRK2 small molecule inhibitor MLi2 (100 nM). **g.** Representative western blots, with quantification in **h.** and **i.**, of lysates from human monocyte-derived macrophages cultured from heathy adult volunteers polarized with human M-CSF for five days and then treated with 1 μg mL^-1^ of human monomer or fibril protein, with or without pretreatment of the LRRK2 small molecule inhibitor MLi2 (100 nM). **j.** Representative immunoblots, and **k., l.**, quantification, of LRRK2 expression and Rab10 phosphorylation from male WT(nTg) mice after intrastriatal injection of 10 μg of α-synuclein fibrils for one month, or saline-injection. Data from graphs show group means as red bars with error bars representing ± SEM from n=3-4 biologically independent experiments or individual mice. Significance is assessed by one-way ANOVA with Tukey’s post hoc test for the indicated group comparisons, except for panels K and L which include significance from 2-tailed t-tests. All human samples used here are derived from PBMCs collected from two healthy volunteer males and one female.

In human CSF1-expanded monocyte-derived macrophages, application of 1 μg per mL of ∼20 nM rod-fibrils (∼642 pM) of human α-synuclein in-solution led to a similar induction of LRRK2 protein expression as in mouse cells (∼2.2 fold, Figure 1G,H) as well as of Rab10 phosphorylation (∼3 fold, Figure 1I). As was seen in mouse macrophages, exposure to MLi2 blocked both Rab10 phosphorylation as well as α-synuclein-induced upregulation of LRRK2 protein expression. Past studies suggested aggregated α-synuclein may stimulate toll-like receptor 4 signaling pathways in myeloid cell activation (Fellner et al., 2013; Hughes et al., 2019; Venezia et al., 2017). However, application of a potent TLR4 inhibitor known to completely block LPS responses failed to block α-synuclein fibril-induction of LRRK2 protein expression and Rab10 phosphorylation (Supplemental Figure 1). These results suggested to us that trace LPS-contamination in α-synuclein fibrils below our limits of endotoxin detection (see Methods) is unlikely to be responsible for LRRK2 induction. Further, the effect on LRRK2 appears specific to fibril preparations since monomeric α-synuclein (with the same degree of potential LPS-trace contamination) lacks LRRK2 induction effect. Together, these results suggest a new connection between increased LRRK2 protein expression and its kinase-activity in macrophages in response to fibrillar forms of α-synuclein.

Emerging evidence suggests immune cell activation follows intracranial injections of α-synuclein fibrils in rodent models, with our past results in rats suggesting recruitment of T-cells and mixed myeloid cells (e.g., CD45^hi^/CD11b+ cells) after α-synuclein fibril injections (Harms et al., 2017a). In cortico-striatal brain lysates from WT (nTg) C57BL/6J mice one-month after intrastriatal injection of 20 nM rod-α-synuclein fibrils, we observed a marked increase in Rab10 phosphorylation with reduced LRRK2 protein (Figure 1J-L). As brain homogenates cannot attribute changes in overall phospho-Rab10 and LRRK2 to one cell type (e.g., different neuronal or astroglial populations that express LRRK2), we investigated tissue sections by immunofluorescence microscopy with the same antibodies to LRRK2. While we were unable to visualize specific signals with the phospho-Rab10 antibody in brain sections, presumably due to insufficient sensitivity of the antibody under conditions of tissue fixation, LRRK2 expression could be observed in rare (compared to LRRK2 expression in neurons) CD68+ (macrosialin) cells (Supplemental Figure 2).

**Figure 2.**
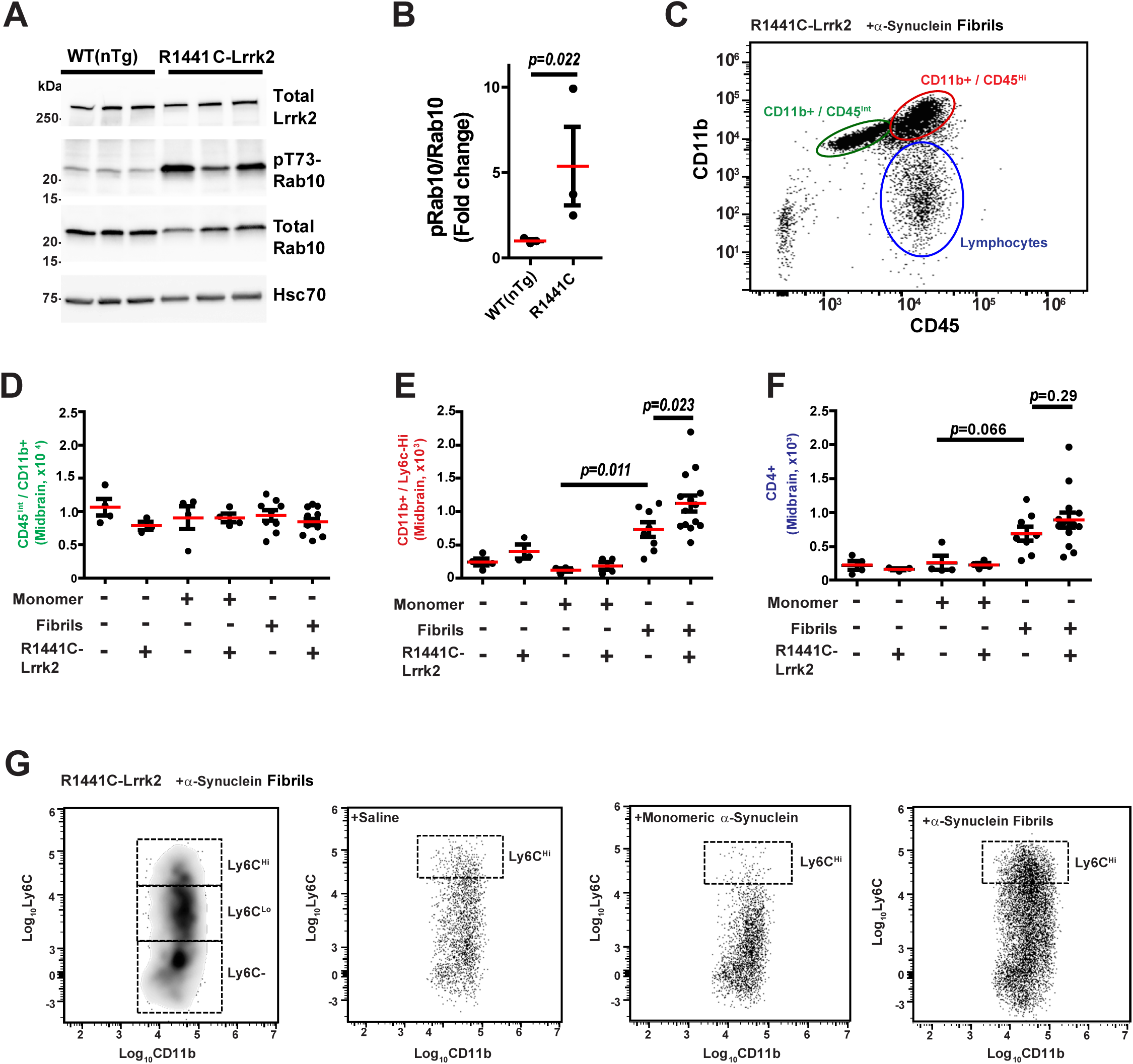
LRRK2 mutation R1441C promotes α-synuclein fibril-induced infiltration of monocytes into mouse brain. **a.** Representative immunoblots, and **b.** quantification, of LRRK2 expression and Rab10 phosphorylation in lysates from dissected frontal cortex and striatum from the brains of male control WT(nTg) mice and homozygous LRRK2-R1441C knock-in mice. Significance is assessed from log_10_ transformed values with a two-tailed t-test, with column graphs showing dots representing individual animals, with red bars indicating group means and error bars representing ± SEM. **c.** Representative scatter plot of live CD45 ^Hi^/CD11b+ cells (MFI) isolated from a mononuclear-enriched Percoll gradient fraction from a selected mouse (male LRRK2-R1441C knock-in) three-days after injection (bilaterally) with 10 μg of α-synuclein fibrils (gating strategy is presented in Supplemental Figure 3). Gates enriched in microglial populations (green), infiltrating and activated myeloid cells (red), and lymphocytes (blue) are indicated. **d.** Dot plots for quantification of the number of CD45^int^/CD11b+ (e.g., microglia). One-way ANOVA is not significant (*p*=0.48). **e.** CD45 ^hi^/CD11b+/Ly6G-/Ly6C^hi^ cells (e.g., classical monocytes), and **f.** CD45^hi^/CD11b-/CD4+ cells (e.g., CD4 T-cells). One-way ANOVA is *p*<0.0002 for both panel e and f. Column graphs show dots representing individual animals, with red bars indicating group means and error bars representing ± SEM from 37 mice (24 females and 13 males, see source data file), 2-3 months in age. *P* values according to Sidak’s multiple comparisons test between the indicated groups. **g.** Representative density plot and scatter plots (MFI) of live CD45^hi^/CD11b+/Ly6G-cells from mouse midbrain cell homogenates three-days after injection (bilaterally) with 10 μg of α-synuclein fibrils, monomer, or saline control. Dashed boxes represent gating strategies marking Ly6C^hi^, Ly6C^lo^, and Ly6C-.

### R1441C-LRRK2 increases pro-inflammatory monocyte responses to α-synuclein fibrils in the mouse brain

We and others have demonstrated LRRK2 control over chemotactic responses in cultured mouse macrophages (Levy et al., 2020; Moehle et al., 2015). To determine whether increased LRRK2 activity is associated with inflammatory responses in the mouse α-synuclein fibril injection model, we employed recently developed congenic (C57BL/6J) R1441C-Lrrk2 knock-in mice that show enhanced phospho-Rab10 levels in mouse-embryonic fibroblasts that is dependent on LRRK2 kinase activity (Steger et al., 2016). The R1441C-LRRK2 mutation results in ∼2-fold increases in phospho-Rab10 in transfected HEK-293T cells (Liu et al., 2017), whereas in these conditions the more common, but less-penetrant, G2019S-LRRK2 mutation has nominal or no significant effect on phospho-Rab10 levels (Liu et al., 2017; Steger et al., 2016). In brain lysates from homozygous R1441C-Lrrk2 mice, we observed an increase (albeit variable) of phospho-Rab10 (Figure 2A,B), consistent with cell lines. We performed flow cytometry on immune cells (focusing on CD45+ cells) from the mouse brain using a gating strategy (Supplemental figure 3) to analyze microglia, infiltrating peripheral monocytes, and lymphocytes (Figure 2C). α-Synuclein fibrils did not induce changes in the number of resident immune cells CD45^int^/CD11b+ cells (i.e., microglia) that always outnumbered monocytes and T-cells in the cell suspensions more than ten-fold (Figure 2D). Interestingly, we observed a sharp increase in the number of CD11b+/Ly6G-/Ly6C^Hi^ monocytes after α-synuclein fibril injection compared to the injection of matched monomer protein or saline injections (Figure 2E). The number of these recruited classical monocytes was further increased over WT(nTg) mice in α-synuclein fibril injected homozygous R1441C-Lrrk2 knock-in mice (Figure 2E, G). There were no genotype effects noted in the injection of matched monomer protein or saline control injections (Figure 2E). α-Synuclein fibrils (but not matched monomer protein) caused an increase in CD4+ T-cells recruited to the brain, although the counts were similar between R1441C-Lrrk2 knock-in mice and WT(nTg) mice (Figure 2F). Overall, these results implicate LRRK2 activity in the recruitment of monocytes to the brain, with the potential for further LRRK2 expression and α-synuclein fibril-induction in the monocytes.

**Figure 3.**
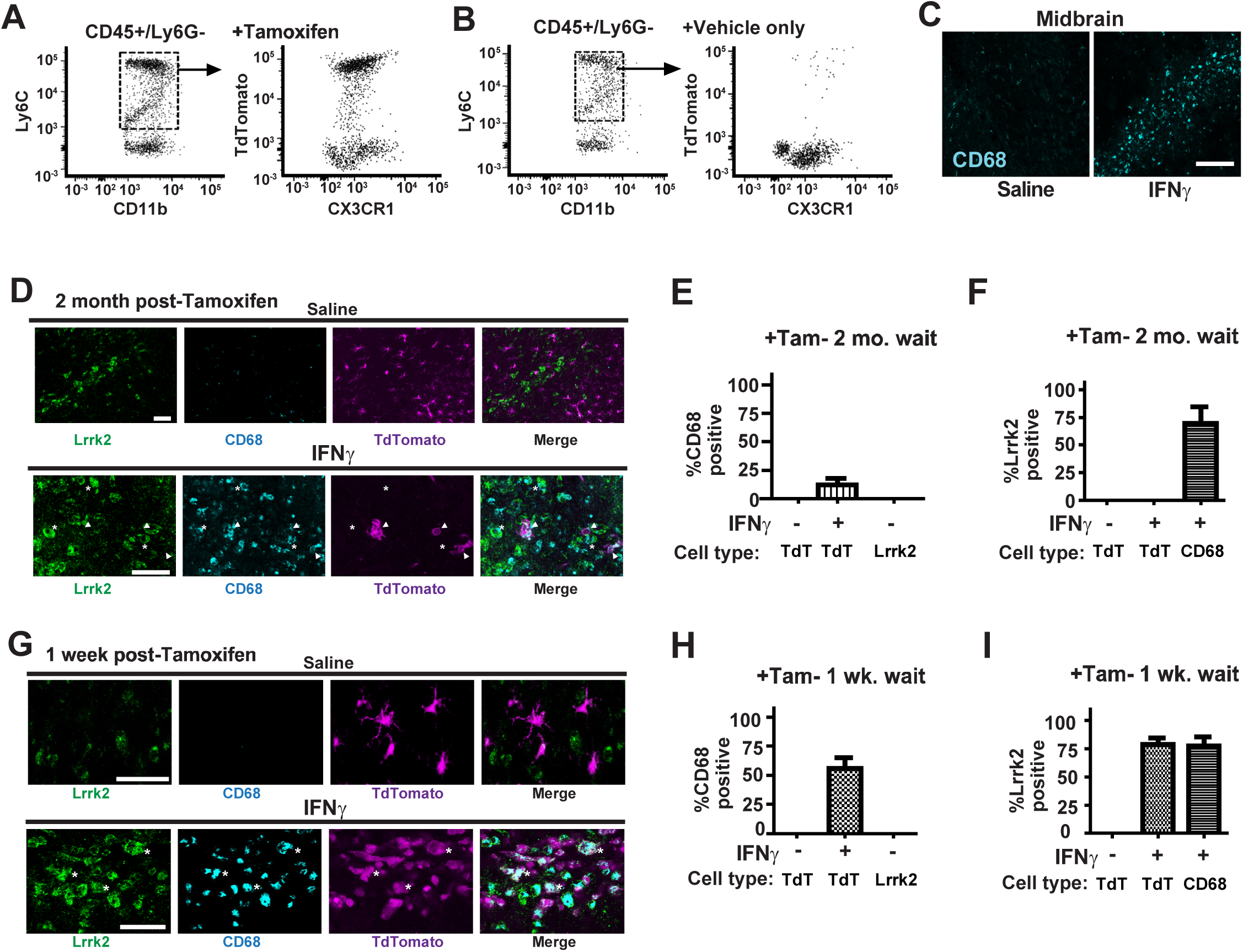
Fate mapping reveals LRRK2+/CD68+ cells are distinct from CNS resident (CX3CR1+) microglia. Double-transgenic mice (CX3CR1-Cre^Ert2^ / Flx-Stop-Flx-tdTomato, see Supplementary Figure 5) were injected in the substantia nigra pars compacta (SNpc) with either saline (as a control) or a combination of 40 ng mouse IFNγ and LPS (10 μg, 10,000 E.U.). 48-hours after injection, mice were evaluated by flow cytometry and immunofluorescence analysis. **a.** Scatter plots of mouse blood from the double-transgenic mice evaluated for Ly6C, CD11b, and tdTomato expression in CD45+/Ly6G-cells by flow cytometry, one-week post-tamoxifen or **b.** vehicle (corn oil) injection. Representative scatter plots (MFI) highlight tdTomato positive expression (from the dashed-line boxes/gates representing Ly6C+ monocytes). Both male and female mice were analyzed per group with similar results (n=3 mice per group; 2 males,1 female each for the vehicle and tamoxifen group). **c.** Immunofluorescence images of a representative coronal section of the mouse SNpc stained for CD68 48 hours post IFNγ injection (cyan). Scale bar is 400 μm. **d.** Representative immunofluorescence images used to quantify co-localization in cells between the expression of LRRK2 (green), CD68 (cyan) and epifluorescence from tdTomato, in cells within the substantia nigra of double transgenic mice, injected with either IFNγ/LPS (sections analyzed 48-hours post injection) or saline control, with IFNγ/LPS injections occurring two-months post-tamoxifen injection. Asterisks denote typical cells recorded as co-positive for CD68 and tdTomato that lack LRRK2 expression, with arrowheads denoting typical cells co-positive for LRRK2 and CD68 (but not tdTomato). **e.** Quantification of cells counted (>100 per group) from at least two sections of SNpc from three male mice each, for CD68 cells co-localized with tdTomato, or LRRK2, and **f.** LRRK2 cells co-localized with tdTomato, or CD68. “+” denotes IFNγ/LPS injection whereas “-” denotes saline control. **g.** Representative immunofluorescence images as in panel d, except IFNγ/LPS injections occurred one-week post-tamoxifen injection and sections evaluated 48 hours later. Asterisks denote typical cells recorded as triple-positive for CD68, tdTomato, and LRRK2 expression. **h.** Quantification of cells counted (>100 per group) from at least three photomicrographs each from three male mice for CD68 cells co-localized with tdTomato, or LRRK2, and **i**., LRRK2 cells co-localized with tdTomato or CD68. Column bars show group means and error bars represent S.E.M from across the images evaluated. All scale bars in panels d and g are 100 μm.

### LRRK2+/CD68+ immune cells recruited to the mouse brain are macrophages distinct from activated microglia

Numerous studies have attributed cell-autonomous LRRK2 and mutant LRRK2-function to microglia both *in vivo* and in cultured microglia-like cells, for example in microglial responses to IFNγ, LPS, and adenosine injections into mouse brain (Choi et al., 2015; Gardet et al., 2010; Kuss et al., 2014; Ma et al., 2016; Moehle et al., 2012; Russo et al., 2019). However, primary microglial cells in culture are known to acquire macrophage-like expression profiles distinct from microglia in the brain (Butovsky et al., 2014), and recent single-cell sequencing efforts have demonstrated low or no *Lrrk2* mRNA expression in resident microglia or resident macrophages in the mouse brain (Supplemental Figure 4). In contrast, in human peripheral blood cells, very high *LRRK2* expression localizes to both CD14+ and CD16+/CD11b+ cells (Supplemental Figure 4 and (Hakimi et al., 2011; Thevenet et al., 2011)). Using newer and more specific antibodies directed to Lrrk2 that we had previously optimized for tissue staining in mouse brain (West et al., 2014), we utilized a fate mapping approach to identify the origins of Lrrk2+ immune cells previously observed as CD68+/IsoB4+ in the brain during neuroinflammation (Daher et al., 2014; Moehle et al., 2012). We created double-transgenic mice combining tamoxifen-inducible Cre expression via a Cx3cr1 promoter with a stop-floxed CAG-promoter driven tdTomato construct (Supplemental Figure 5, and (Parkhurst et al., 2013)). Upon tamoxifen administration, CX3CR1+ cells (e.g., microglia) and their progeny were irreversibly labelled by TdTomato, in contrast to blood monocytes where negligible numbers of CD11b+/Ly6G-/Ly6C^Hi^ monocytes were tdTomato positive one-month after tamoxifen injection, or with vehicle-only injection (Figure 3 and Supplementary Figure 5). However, one-week post tamoxifen injection, we found that most CD11b+/Ly6G-/Ly6C^Hi^ monocytes in blood were also tdTomato positive, with tdTomato expression being dependent on tamoxifen (Figure 3A,B). These results are consistent with the longer half-life associated with microglia (>3-months (Fuger et al., 2017)) compared to the shorter half-life for Ly6C+ mouse blood monocytes (<7 days, (Yona et al., 2013)) where no blood cell labeling can be detected in as little as one-month post-tamoxifen injection.

**Figure 4.**
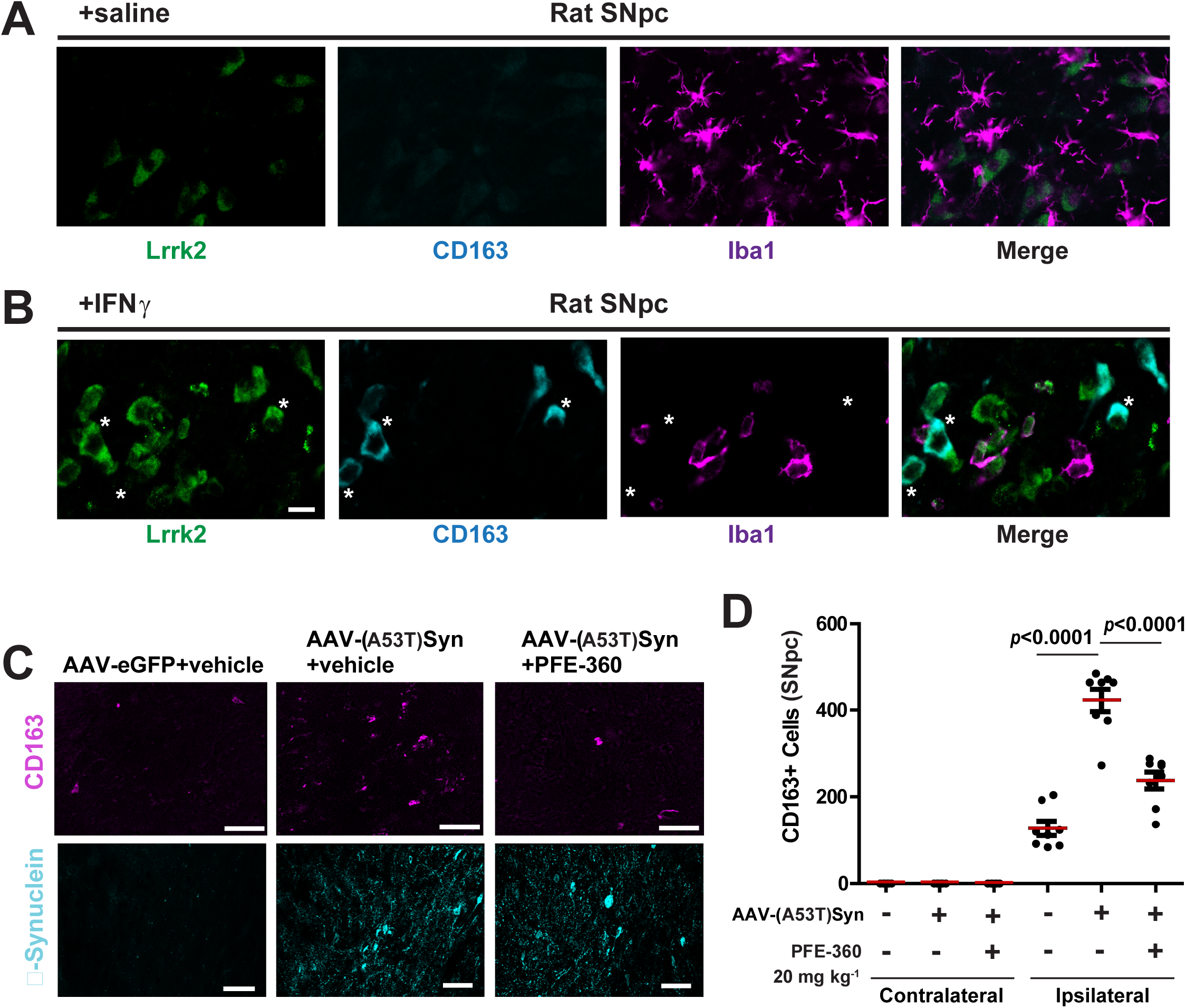
LRRK2 co-expresses with CD163 macrophages in the rat brain. **a.** Representative confocal images of coronal sections in the substantia nigra pars compacta (SNpc) from a male Sprague Dawley rat injected in the SNpc with either saline as a control, or **b.**, a combination of 40 ng mouse IFNγ and LPS (10 μg, 10,000 E.U.). 48-hours post-injection, tissue was evaluated by immunofluorescence analysis with antibodies to LRRK2 (green), CD163 (cyan), and Iba1 (magenta). Representative high-magnification immunofluorescence in coronal sections in the rat SNpc are shown, scale bars are 10 μm. Asterisks denote nearby cells that are LRRK2+/CD163+ and Iba1(-)/weak. CD163+ and Iba1(+)/strong cells in blood vessels were LRRK2 negative (see Supplemental Figure 7). **c**. Male Sprague Dawley rats were unilaterally injected into the SNpc with 6.8×10^9^ viral genomes (rAAV2/1) encoding eGFP as a control, or human A53T-α-Synuclein, and brains collected four weeks after rAAV2/1 injection. Representative immunofluorescence images are show with CD163 staining (magenta) and nitrosylated/misfolded α-synuclein antibody (Syn514 monoclonal antibody, cyan), scale bars are 200 μm. **d.** Stereological counts (rare-object counting protocol) of CD163+ cells in the injected (ipsilateral) and contralateral parenchymal SNpc. “+” indicates human A53T-α-Synuclein virus and “-” indicates eGFP virus, and “+” indicates PFE-360 treatment, expected to completely inhibit LRRK2 kinase activity over 24 hours (see Supplemental figure 7), whereas “-” indicates vehicle control treatment. Red lines depict group means, and error bars show ± SEM, with dots representing assessments of estimated total counts through the SNpc of individual male rats (n=8-10 per group, see Source data). Significance is assessed by one-way ANOVA (*p*<0.0001), and indicated group mean significance is according to Sidak’s multiple comparisons test.

**Figure 5.**
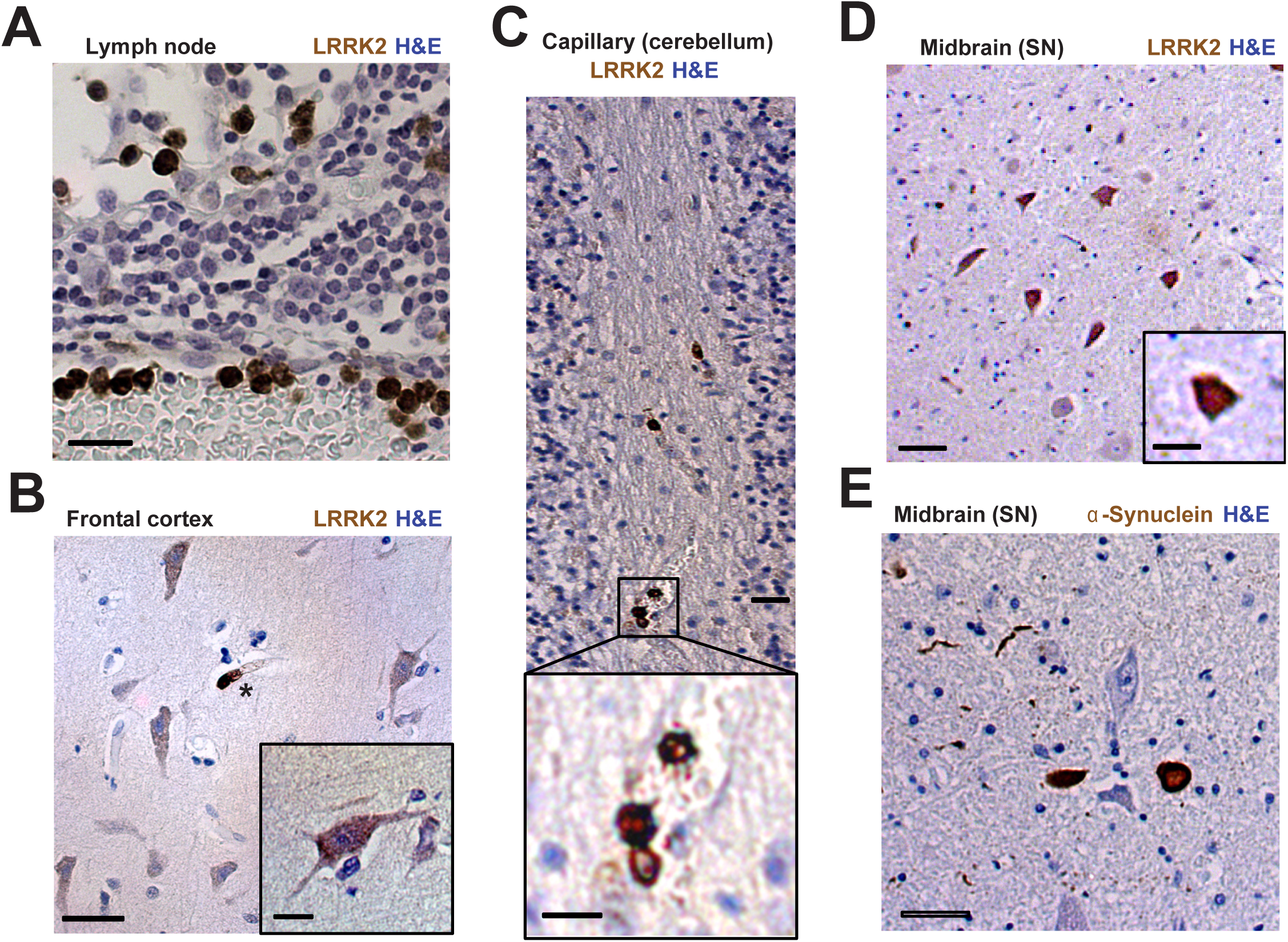
LRRK2 expression in macrophage-like cells in PD midbrain with α-synuclein pathology. **a.** Robust cytoplasmic LRRK2 reactivity (brown) in granulocytes and monocytes in the mesenteric lymph node, with hematoxylin counterstain (blue). **b.** Representative LRRK2 reactivity (total LRRK2 protein) in the brain of a patient with advanced, typical Parkinson’s disease. Shown are sections from frontal cortex with LRRK2 reactivity highlighting pyramidal neurons (including in the higher magnification inset) and an intensely stained myeloid cell within a nearby capillary (denoted by an asterisk). **c.** Prominent LRRK2 expression in intravascular cells of a white matter tract in the cerebellum. High-magnification inset shows a trio of amoeboid-shaped cells with prominent cytoplasmic LRRK2 expression. **d.** LRRK2 reactivity in cells (<20 μm) with macrophage-like morphological features in the substantia nigra pars compacta stained with **e.** anti-α-synuclein (pS129) in serial sections, highlighting two prominently stained aggregates (axonal spheroids or former intracellular Lewy bodies) as well as numerous Lewy neurites. Scale bars are 40 μm for main photomicrographs and 20 μm for high-magnification insets. Images are representative of three idiopathic PD brain samples evaluated.

In mice with tdTomato expression in microglia (2-month wait after tamoxifen injection), or with tdTomato expression in both microglia and blood monocytes (1-week wait after tamoxifen injection), we injected a combination of mouse IFNγ and LPS intracranially to acutely induce blood-brain barrier breakdown and monocyte extravasation to explore LRRK2 expression in different immune cells recruited to the brain. Two-days after intracranial injection, numerous CD68+ myeloid cells, representing different classes of microglia, macrophages, and other cell types, were observed distributed across the substantia nigra (Figure 3C). In the 2-month wait after injection of tamoxifen to eliminate any peripheral tdTomato labeling 48-hours post LPS/IFNγ injection, there were no instances of a cell with both tdTomato and LRRK2 expression (Figure 3D-F). In contrast, analysis of mice 1-week post tamoxifen, 48 hours after LPS/IFNγ injection, revealed a strong co-localization (∼80% of all CD68 cells) between LRRK2 expression in CD68 cells (Figure 3G-I). Together, these results suggest that LRRK2-expressing immune cells in the mouse brain must be distinct from long-lived brain resident microglia and macrophages and their progeny in LPS/IFNγ induced neuroinflammation.

As our flow cytometry analysis suggests Cx3cr1-Cre efficiently genetically labels peripheral LRRK2-expressing cells recruited to the brain shortly (i.e., 2-days) after intracranial LPS/IFNγ injections, we tested whether a Cx3cr1^WT/eGFP^/CCR2^WT/RFP^ double knock-in mouse, traditionally used to separate brain-resident cells from CCR2-expressing peripheral monocytes (Jung et al., 2000; Saederup et al., 2010), may help rapidly differentiate LRRK2 expressing immune cells in the brain in neuroinflammation. However, the results were ambiguous (Supplemental Figure 6), with LRRK2+/ CD68+ double-positive amoeboid cells weakly and variably positive for both RFP (CCR2) and eGFP (Cx3cr1), despite an abundance of both eGFP-only and RFP-only cells in the brain due to the LPS/IFNγ injection. Taken together, these results suggested that murine *Lrrk2* expression may be upregulated, while CCR2 and Cx3cr1 levels may be downregulated, upon IFNγ/LPS-induced immune cell extravasation into the brain parenchyma.

**Figure 6.**
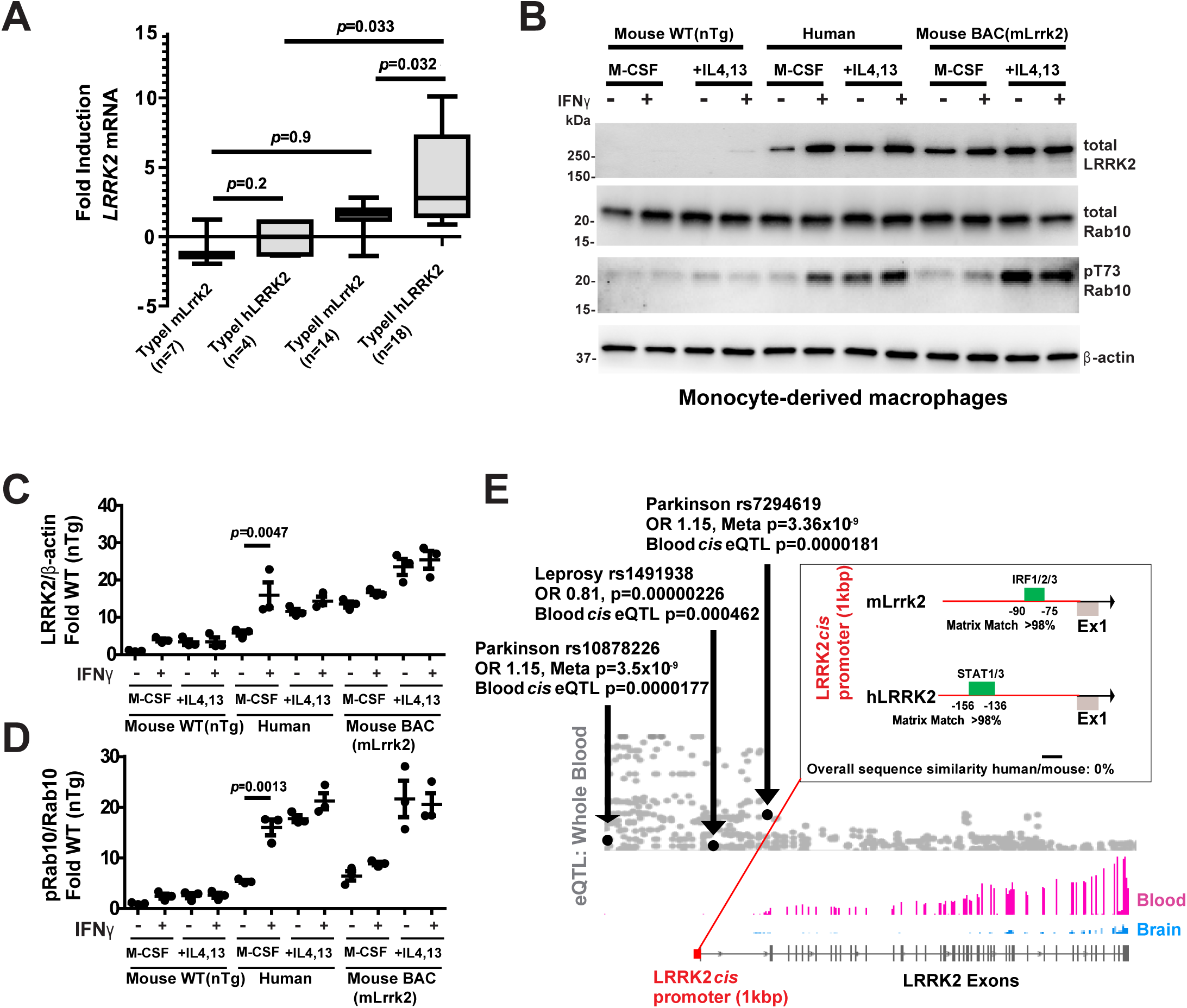
LRRK2 activation in type II interferon signaling in human but not mouse cells. **a.** Analysis of *LRRK2* mRNA induction from 43 experiments from immune cells (n indicated in each group) as indexed in the Interferome database of IFN-inducible genes (interferome.org). One way ANOVA is *p*<0.0003, with Tukey’s post-hoc test results for the indicated group mean comparisons. **b.** Representative immunoblots of lysates from bone marrow-derived macrophages from WT(nTg) control mice, human monocyte-derived macrophages, and bone marrow-derived macrophages from mouse-mBac WT-LRRK2 over-expressing mice. Cells were polarized with M-CSF in culture for five days and then further treated with or without a combination of IL4 and IL13 for an additional two days, and then treated for 48 hours with IFNγ (20 ng mL^-1^) as indicated. **c.** Immunoblot quantification of LRRK2 levels normalized to β-actin, and **d.**, phospho-Rab10 normalized to total Rab10, calculated from three independent cultures from three male mice each and from three healthy human volunteers (see Methods). Data are expressed as fold change relative to the WT(nTg) control mice without IFNγ stimulation (first lane of blots in panel b.). Lines in graphs show group means with error bars depicting ± SEM. Significance is assessed by one-way ANOVA with Tukey’s post hoc test for the indicated group in the effect of IFNγ stimulation. **e.** Relative expression (GTEx IGV Browser) of the 51 exons in human *LRRK2* are depicted in blood (magenta) versus brain (cyan) expression, with eQTLs plotted and the positions of selected GWAS PD- (Nalls et al., 2014) and leprosy-associated *LRRK2* promoter variants (Fava et al., 2016; Zhang et al., 2009) indicated. *P* values associated with blood eQTLs are adapted from the blood eQTL browser (genenetwork.nl/bloodeqtlbrowser/, (Westra et al., 2013). Finally, IFN-associated transcription factor binding sites with respect to the mouse and human *LRRK2* transcription start site are indicated, as identified in the Interferome database of IFN-inducible genes (interferome.org).

### LRRK2 mediates the recruitment of CD163+ macrophages in rats after rAAV2-A53T-α-synuclein transduction

A monoclonal antibody to the rat hemoglobin-scavenger receptor (CD163, clone ED2) has been proposed to demarcate mature and polarized monocyte-derived macrophages and perivascular macrophages in rat brain (Polfliet et al., 2006). To determine whether LRRK2 expression co-localizes with CD163 macrophages in the rat brain, and is involved in their recruitment, we first analyzed LRRK2 expression in parenchymal CD163 cells after IFNγ/LPS injection. In saline injection conditions (Figure 4A), LRRK2 expression was confined to neurons, consistent with previous immunohistochemical localization of LRRK2 in rat brain (West et al., 2014). However, with IFNγ/LPS-induced neuroinflammation, all detectable CD163+ cells were co-positive with LRRK2 expression in the brain parenchyma (Figure 4B), although LRRK2 expression was not detected in CD163+ perivascular macrophages or Iba1+ cells along vessel walls from PBS-perfused brains (Supplemental Figure 7A). The co-localization between LRRK2 and CD163, a marker of monocyte-derived macrophages in rats, suggest LRRK2+ immune cells in the brain can originate from blood cells.

**Figure 7.**
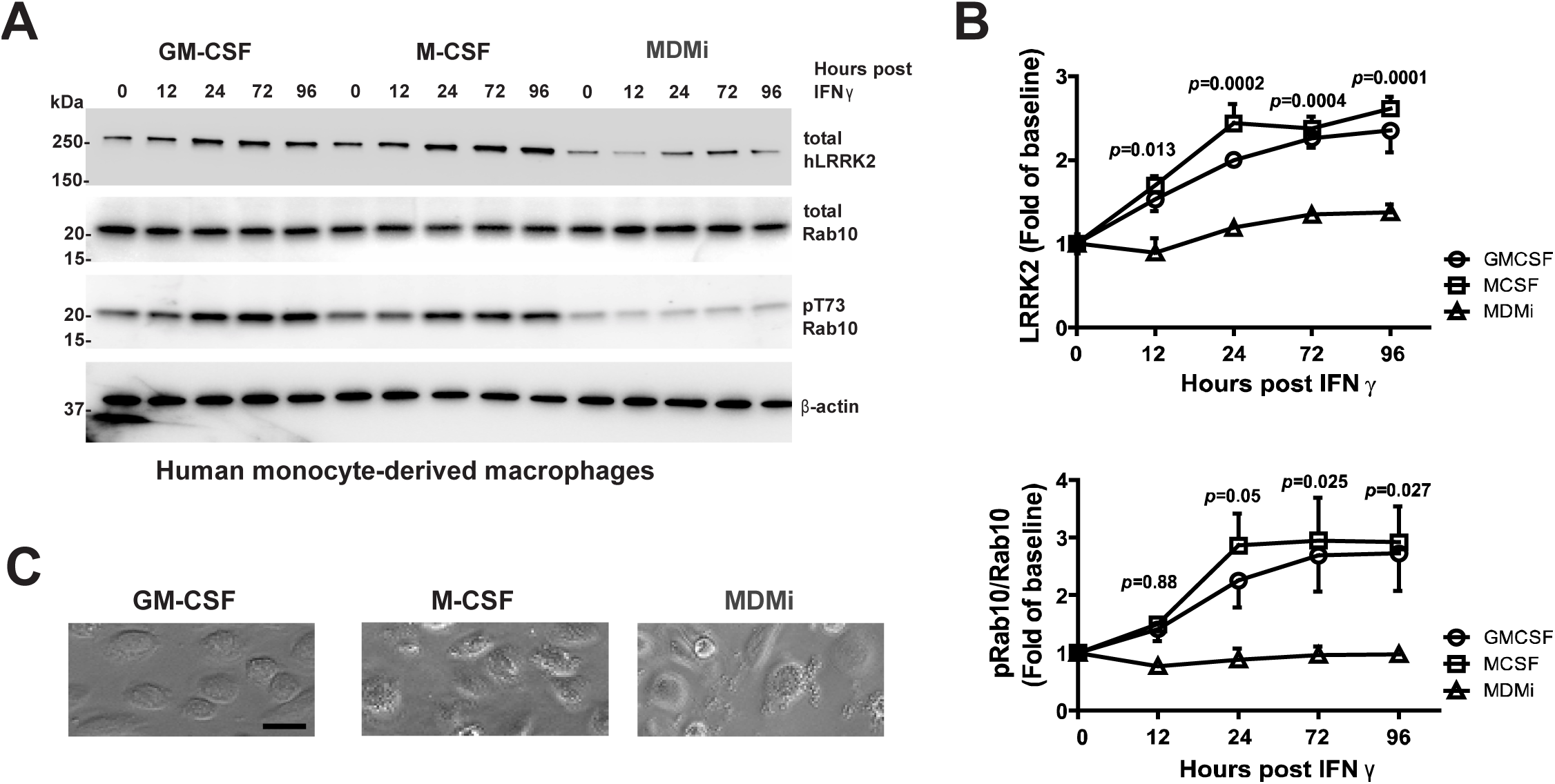
Human monocyte polarization to microglia-like cells suppresses late and sustained IFNγ-LRRK2 induction in macrophages and dendritic-like primary cells. **a.** Representative immunoblots from lysates of human peripheral monocytes polarized to dendritic-like cell (GM-CSF, 5 days), macrophage (M-CSF, 5 days) and microglia-like cells (GM-CSF, M-CSF, NGFβ, CCL2, and IL-34, 10 days). Cells were treated with human IFNγ (20 ng mL^-1^) and collected at the indicated time points post IFNγ treatment. **b.** Quantification of LRRK2 levels, normalized to β-actin, and phospho-Rab10 levels normalized to total Rab10 levels, calculated as fold change relative to the 0-hour time point (i.e, no IFNγ) of each cell type. Data are calculated from experiments from cells from two healthy male volunteers, with cells plated in triplicate experiments each, with group means shown with error bars as ± SEM. No significant differences were observed between the GM-CSF and M-CSF treated groups at any time point. Significance in GM-CSF/M-CSF groups compared to MDMi groups is assessed by one-way ANOVA (p<0.0001 for both LRRK2 and phospho-Rab) with Tukey’s post hoc test results at the indicated time point representing comparisons of GM-CSF and M-CSF responses with MDMi responses. **c.** Representative bright field images of cultured cells before IFNγ stimulation, with scale bar at 50 µm. M-CSF:macrophage colony-stimulating factor; GM-CSF:granulocyte-macrophage colony-stimulating factor; MDMi: monocyte-derived microglia-like cells.

We next aimed to test whether LRRK2 kinase activity mediates CD163+ cell recruitment in a rat model of PD. Previously, we demonstrated that LRRK2 knockout rats have fewer CD68+ cells recruited to the brain compared to WT(nTg) rats after rAAV2-α-synuclein viral injections (Daher et al., 2014). Some CD68+ cells that accumulate in rat brain in this model were suspected to be monocyte-derived macrophages based on morphological analyses (Sanchez-Guajardo et al., 2010). Using the LRRK2 kinase inhibitor PFE-360 (Kelly et al., 2018), twice daily oral gavage doses resulted in the complete ablation of LRRK2 kinase activity, as monitored by pS935-LRRK2 levels as described (Supplemental Figure 7, and (Kelly et al., 2018)). PFE-360 has improved bioavailability compared to the closely related molecule PFE-475 we have used in the past, allowing lower quantities of drug to achieve similar inhibition profiles (Daher et al., 2015; Kelly et al., 2018). In an early time point one-month post rAAV2-A53T-α-synuclein viral injection, the number of CD163+ cells in the midbrain parenchyma were counted using unbiased stereology. We found that the number of CD163+ cells recruited to the ipsilateral injection site were markedly reduced with LRRK2 kinase inhibition (Figure 4C,D). No CD163+ cells in the uninjected side of the brain were found. Notably, these cells are rare, measuring in the hundreds across the rAAV2-A53T-α-synuclein injected SNpc (Figure 4D). These results suggested to us that LRRK2 upregulation in monocyte-derived macrophages may be conserved from mice to rats in immunological responses to abnormal α-synuclein, in this case in response to A53T-human-α-synuclein over-expression in the rat midbrain.

### LRRK2 expression in human monocytes and the brain in PD

Whether monocyte-derived macrophages are present in susceptible brain parenchyma of PD brain and other neurodegenerative disease remains controversial, principally due to a lack of antibodies that convincingly separate the different lineages of myeloid cells in the brain. To identify an immunohistochemical method for LRRK2 detection in formalin-fixed sections, we first probed LRRK2 expression using antibodies previously validated in Lrrk2 knockout cells for immunoreactivity in mesenteric human lymph nodes that are enriched with monocytes. Anti-human LRRK2 antibody, HL-2, previously described (Berger et al., 2010; Hakimi et al., 2011), was effective in highlighting abundant LRRK2+ amoeboid-shaped cells with prominent cytoplasmic LRRK2 expression, consistent with LRRK2 reactivity previously characterized in monocytes and macrophages from mice and rats (Figure 5A). Brain sections from three idiopathic PD cases and three control brains without neurodegenerative disease were evaluated with HL-2 antibodies. In neocortical tissue sections from idiopathic PD with sparse or no α-synuclein pathology (*LRRK2*-mutation negative), typical LRRK2 expression in pyramidal neurons could be resolved (Davies et al., 2013; Higashi et al., 2007). Intensely labeled, small (10 μm or less) cells could be occasionally identified in microcapillaries within the cortex (Figure 5B). LRRK2+ amoeboid cells could also be identified in larger capillaries in transverse sections of white matter tracts (Figure 5C). In sections of PD midbrains with Lewy pathology, clusters of small LRRK2+ cells could be identified with flatter and polarized morphologies than observed for the LRRK2+ cells in capillaries (Figure 5D). In adjacent serial sections of the same midbrains processed in parallel for α-synuclein reactivity, we detected numerous intracellular Lewy-type inclusions (Figure 5E) as well as extracellular aggregates. Neither LRRK2+ small cells or Lewy-type inclusions were detected in brain parenchyma in control brain sections. Thus, rare LRRK2+ cells in the human PD midbrain were morphologically similar to LRRK2+ monocyte-derived macrophages labeled in mouse midbrain and CD163+ macrophages labeled in rats with α-synuclein-induced neurodegeneration.

### LRRK2 activation in type II interferon signaling in human but not mouse macrophages

Abnormal or aggregated α-synuclein protein stimulates immune cells through a number of different pathways including stimulation of IFNγ (Refolo and Stefanova, 2019; Sarkar et al., 2020; Vieira et al., 2015). Previous reports had suggested LRRK2 expression induced during type II interferon (IFN) responses associated with IFNγ (Gardet et al., 2010; Kuss et al., 2014). Aggregated α-synuclein protein in PD may stimulate the secretion of pro-inflammatory molecules in microglia that can include IFNγ (Grozdanov et al., 2019; Harms et al., 2017b; Hughes et al., 2019; Wang et al., 2019), and IFNγ is a major pro-inflammatory cytokine in Crohn’s disease and macrophage activation in mycobacteria responses (Cooper and Khader, 2008; Strober et al., 2010). However, a meta-analysis of 43 previous gene expression studies in immune cells, as indexed in the Interferome database of IFN-regulated genes, indicated only a nominal or no increase at all for *Lrrk2* mRNA in IFNγ stimulation of immune cells (Rusinova et al., 2013). In contrast, *LRRK2* mRNA was reliably induced over baseline levels with IFNγ stimulation in human immune cells (Figure 6A).

To evaluate phospho-Rab10 levels and LRRK2 expression induction in monocyte-derived macrophages after IFNγ stimulation, we cultured cells from monocytes classically (M-CSF) polarized and alternatively activated (IL4, IL13-stimulated), with or without IFNγ stimulation, from WT(nTg) mice versus that of healthy human controls. LRRK2 expression in mouse macrophages, at baseline and after IFNγ stimulation, was an order of magnitude less than the expression in human macrophages (normalized to β-actin and total protein). A large increase was detected in both LRRK2 expression and LRRK2 kinase activity with IFNγ stimulation in human cells, with more nominal differences observed in mouse WT(nTg) cells (Figure 6B-D). IFNγ stimulation in classical polarized cells increased LRRK2-driven Rab10 phosphorylation comparable to that observed in alternatively activated cells which had elevated LRRK2 kinase activity without IFNγ stimulation. To ensure that a lack of LRRK2 induction with IFNγ stimulation in mouse monocyte-derived macrophages is not because of very low levels of endogenous Lrrk2 expression inherent to WT(nTg) mouse cells, we generated mice with ∼30 copies of a BAC transgene encoding the WT mouse *Lrrk2* gene to boost expression to that of human macrophages (Figure 6B-D). However, IFNγ stimulation still had little effect on LRRK2 induction or kinase activity in BAC-transgenic mouse LRRK2 cells, in either classical or alternatively activated cells. Notably, the level of phospho-Rab10 and LRRK2 expression were comparable between BAC-transgenic mouse cells and human cells in alternatively activated (+IL-4,13) cells, with comparable LRRK2 kinase activity in alternatively activated cells in both human and BAC-transgenic mouse cells.

To gain further insight into the difference between mouse and human cells with respect to IFN responses and LRRK2 expression, transcription factor binding prediction based on *in silico* modeling revealed an IRF1/2/3 binding site in the mouse *cis Lrrk2* promoter (<1kbp upstream of the *LRRK2* transcription start site), whereas a STAT1/3 binding site is found in the human *cis LRRK2* promoter (Figure 6E). Of note, neither human or mouse promoter sequences contained the canonical pairs of IRF and STAT bindings sites usually found in IFN-regulated genes. While LRRK2 amino acid sequence is highly conserved in mammals (West, 2015; West et al., 2014), overall there is striking divergence between the human and mouse *LRRK2 cis*-promoter sequence, with no significant sequence similarity or transcription factor binding sites between species (Figure 6E). Further interrogation of the human *LRRK2* promoter revealed genome-wide associated variants in *LRRK2* for PD and leprosy (Hansen’s disease) are expression quantitative trait loci (eQTLs) in blood but not in brain tissue (Figure 6E).

We next determined the timing of IFNγ induction of LRRK2 expression and activity in human cells, and how different cell polarizations affect the relationship between IFNγ and LRRK2. Both M-CSF polarized (e.g., macrophage-like) and GM-CSF (e.g., dendritic-like) cells demonstrated very late LRRK2 responsiveness to IFNγ stimulation, with peak activity observed 24-hours post IFNγ that sustained for days (Figure 7A,B). Phopsho-Rab10 induction mirrored that of LRRK2 protein expression induction without significant changes in total Rab10 protein. The LRRK2 kinase inhibitor MLi2 completely blocked phospho-Rab10 levels at baseline and after IFNγ stimulation of both M-CSF and GM-CSF cell cultures (Supplemental Figure 8). No significant differences in LRRK2 expression or induction of activity towards phospho-Rab10 were noted between M-CSF and GM-CSF cells. Recently, protocols to polarize monocyte-derived cells towards microglia-like phenotypes have been identified (Ryan et al., 2017). Based on our findings from above, we wondered whether these cells, cultured from the same subjects at the same time as the M-CSF and GM-CSF cells, might actually diminish LRRK2 expression and reduce IFNγ induction of LRRK2 activity. In monocyte-derived microglia-like (MDMi) cells, LRRK2 expression and phospho-Rab10 (but not total Rab10) levels were lower at baseline and showed no response after IFNγ stimulation (Figure 7A,B). Despite the biochemical differences between the three polarized states of immune cells studied here (GM-CSF, M-CSF, MDMi), morphologically, these cells were highly similar in appearance (Figure 7C).

## Discussion

The present study focused on LRRK2 protein expression and activity in relation to pathogenic changes in α-synuclein. Novel conclusions center on three experimental lines of evidence. First, α-synuclein fibrils potently stimulate LRRK2 activity in both mouse and human monocyte-derived macrophages. Notably, LRRK2 expression and kinase activity toward the LRRK2 kinase substrate Rab10 is at least an order-of-magnitude higher in human cells than in mouse cells, whereas total Rab10 protein is equivalently expressed in the cells studied here. Second, LRRK2 expression and kinase activity is highly suppressed in microglial populations but robust in peripheral monocyte-derived cells that might be recruited to the brain in response to abnormal α-synuclein and neuroinflammation. Third, we provide additional support in a mouse and rat model of PD for the hypothesis that LRRK2 expression and kinase activity may regulate chemotactic responses in monocyte-derived macrophage recruitment to the brain. Taken together, these results provide evidence that LRRK2 may drive neuroinflammatory responses in part through control of peripheral monocytes recruited to the brain in disease.

Therapeutics targeting LRRK2, including anti-sense oligonucleotides and small molecule LRRK2 kinase inhibitors, have recently advanced through safety trials in the clinic (e.g., ClinicalTrials.gov NCT03976349 and NCT04056689) in preparation for treating PD patients with *LRRK2* mutations as well as idiopathic PD (West, 2017). A better mechanistic understanding of how LRRK2 kinase activity ties with α-synuclein aggregation that pathologically defines PD may increase the probability of identifying successful clinical approaches. As opposed to α-synuclein, LRRK2 does not aggregate in disease and is expressed in both neurons and cells in the immune system. Pathogenic *LRRK2* mutations appear to uniformly hyperactivate LRRK2 kinase activity in cells that express LRRK2 protein (West, 2017; West and Cookson, 2016). While carriers of pathogenic *LRRK2* mutations with PD cannot be clinically distinguished from idiopathic PD, over a quarter of *LRRK2* mutation carriers with PD lack any detectable α-synuclein pathology in their brains (Kalia et al., 2015). These results lead some to speculate that factors other than α-synuclein, particularly those that may lead to deleterious pro-inflammatory responses in the brain, occur downstream of mutant *LRRK2* in the demise of vulnerable neuronal populations that give rise to pleomorphic phenotypes of parkinsonism (Kalia et al., 2015; Rajput et al., 2006; West, 2017; Zimprich et al., 2004). Indeed, IFNγ expression (and LPS injection) alone into the mouse midbrain leads to selective dopaminergic neurodegeneration (Chakrabarty et al., 2011; Mount et al., 2007), while dopamine neurons in the SNpc are well known to be selectively vulnerable to α-synuclein over-expression or aggregation. Recent evidence in mouse and rat models suggests *LRRK2* knockout diminishes pro-inflammatory responses to a variety of exogenous stimuli that include LPS, IFNγ, and adenosine (Dwyer et al., 2020; Hakimi et al., 2011; Kozina et al., 2018; Schapansky et al., 2015). Therefore, anti-inflammatory effects may represent important endpoints to study in clinical evaluation of successful LRRK2-targeting therapeutics, if such effects and cells underlying the effects were better understood.

Dozens of studies, including some from our group, have identified intrinsic LRRK2 function in microglial-like cells that include primary cultured microglia and mouse BV-2 cell lines. However, microglia in culture are known to diverge in transcription profiles towards macrophage signatures distinct from microglia in the brain (Butovsky et al., 2014). Emerging single-cell RNA sequencing profiles from mouse brain fail to identify significant LRRK2 transcript in resident immune cells, leading us to speculate here that LRRK2+ immune cells may be distinct from resident innate immune cells in the brain. Single-cell sequencing data may provide an explanation in our past studies with LPS or α-synuclein over-expression where LRRK2 robustly localizes with CD68 but fails to co-localize in cells with high Iba1 expression (Daher et al., 2014; Moehle et al., 2012). However, CD68 expression is inducible in several lineages of cells in the innate immune system and typically associates with pro-inflammatory (e.g., IFNγ) responses (Pillai et al., 2009). Thus, directed studies were required to explore LRRK2 expression lineage in immune cells as well as LRRK2 kinase activity towards the LRRK2 substrate Rab10 in the context of pro-inflammatory stimuli.

Multiple approaches in mice, some invasive and others genetic-based, have been shown to effectively map lineages of innate immune cells important in disease. Bone marrow chimerism is an effective approach commonly used to differentiate microglia from monocyte-derived macrophages. However, the irradiation used may cause the proliferation of different subpopulations of myeloid cells that makes tracing studies difficult to interpret. To circumvent this problem, parabiosis is an effective alternative, although the low and variable yield of chimerism typical for parabiosis methodologies may introduce confounding variables difficult to control in determining ratios of LRRK2-expressing cells in CD68 cell populations detectable with our immunohistochemistry methods. Similarly, rates of low chimerism are also typical for adoptive transfer methods. We reasoned that Cx3cr1-Cre^ERT2^ knock-in provide an effective non-invasive and unbiased tracking method, harnessing the innate difference in longevity between microglia and the short lifespan of circulating monocytes post-tamoxifen exposures. Empirical measures supported this approach, as tamoxifen induction led to labeling of nearly all Ly6C+ monocytes in the mice. The life-span of the labeled cells in the blood was short lived, as expected, undetectable after one month. This approach was unambiguous in demonstrating LRRK2-expressing CD68+ cells are distinct from microglia, and other long-lived resident immune cells, and their activated progeny in acute IFNγ stimulation. We offer further support for our conclusions with the observation of robust LRRK2 co-localization with the monocyte-derived macrophage marker rat CD163 in brain parenchymal tissue, but not in Iba1+ cells. Although the data suggest we captured a majority of LRRK2+ immune cells in the brain with our approaches, there remains the possibility of important LRRK2 function in immune cells with lower LRRK2 expression not-detectable with our confocal analysis, or in immune cells that are rare or not recruited at the time points evaluated in our α-synuclein and IFNγ/LPS paradigms.

While we have been unfortunately unable to detect LRRK2 expression or phospho-Rab10 expression via flow cytometry analysis of mouse cells, presumably due to the relatively low levels of LRRK2 in murine Ly6C+ monocytes compared to human CD14+ monocytes, or non-optimized procedures, three independent studies demonstrated LRRK2 upregulation in CD14+ monocytes purified from idiopathic PD patients (Bliederhaeuser et al., 2016; Cook et al., 2017; Kim et al., 2018b). Our data suggest that LRRK2 is expressed much higher in human cells, and boosting LRRK2-expression using BAC-transgenic technology to equal that of LRRK2 expression in human cells as described here presents a promising approach to develop and integrate powerful cytometry approaches coupled with emerging single-cell transcriptomic analyses to further decipher LRRK2+ immune cells in the brain. Whether equivalent monocyte-derived macrophages invade the brain in PD remains controversial. Unfortunately, the same anti-rat CD163 antibody clone used to delineate monocyte-derived macrophages in rats appears to uniformly cross-react with microglia cells in the human brain (Pey et al., 2014).

Presently, there are no known antibodies successful in distinguishing monocyte-derived macrophages from other myeloid cells in human brain, nor are the potential function of these cells in disease specifically explored in past studies. We present data in PD brain tissue that anti-LRRK2 antibodies will be promising candidates to explore more broadly in human PD tissue and other diseases of neuroinflammation, especially if ongoing single-cell sequencing efforts through the human innate immune cell compartment support our hypothesis that LRRK2 expression effectively demarcates monocyte-derived macrophage transcriptional signatures in the brain. However, it will be challenging to attribute specific lineages to transient innate immune cell signatures found in human brain tissue. Further complicating the approach, subpopulations of T-cells (both CD4 and CD8) clearly occur in brain parenchymal tissue in PD (McGeer et al., 1988). Our data are consistent with others in that LRRK2-linked PD GWAS disease susceptibility variants present as strong eQTLs in blood but not brain tissue. An alternative similarly attractive hypothesis advanced by others suggests that LRRK2 genetic variants regulate monocyte responses in the periphery that indirectly lead to differences in soluble factors that can cross the blood-brain barrier, without these cells necessarily infiltrating the brain to influence disease responses (Kozina et al., 2018). With either hypothesis, our results and others firmly place LRRK2 expression and inducible host activity in peripheral myeloid cells during the responses to diseases of the nervous system including virally induced encephalitis (Shutinoski et al., 2019).

Although many different abnormal conformations and post-translational modifications of α-synuclein have been associated with neurotoxicity and PD, in the last ten years, recombinant α-synuclein short-fibrils have gained particular notoriety for their ability to template new fibril formation from endogenous α-synuclein expression in neurons (Luk et al., 2012; Volpicelli-Daley et al., 2011). Our results in rat brain demonstrate that these fibrils are not inert with respect to neuroinflammation, but demonstrate that intracranial injections of fibrils (but not monomeric protein) result in significant recruitment of CD45^hi^/CD11b+ cells and T-cells to the rat brain (Harms et al., 2017a). Experiments *ex vivo* in human monocytes demonstrate that α-synuclein rod-fibrils specifically result in pro-inflammatory responses at very low concentrations that includes the production and release of IL-6 (Grozdanov et al., 2019). We find that similar preparations of fibrils also induce LRRK2 expression and kinase activity in monocyte-derived macrophages. While LRRK2 expression and activity affect α-synuclein fibrillation in neurons in a multitude of models (Bae et al., 2018; Bieri et al., 2019; Lin et al., 2009; Volpicelli-Daley et al., 2016; Zhao et al., 2017), our data here add another layer of complexity to the interaction between LRRK2 and pathogenic α-synuclein, namely in inducible LRRK2 activation by immune cells, a property conserved from mice to humans.

We provide the first flow-cytometric analysis of α-synuclein fibril-injected mouse brain to demonstrate robust recruitment of both Ly6C+ monocytes as well as T-cells. In evaluation of the recently developed R1441C knock-in mice, a pathogenic LRRK2 mutation well known to cause Rab10 hyperphosphorylation, the number of monocytes recruited to the brain are markedly increased. These data may further implicate intrinsic LRRK2 expression and kinase activity in chemotactic responses in monocytes, and possible T-cell responses. An alternative hypothesis advanced by others suggests LRRK2 primary action in human monocytes may involve maturation and survival of monocytes in tissues after IFNγ stimulation (Thevenet et al., 2011). Although IFNγ / LPS exposures of mice and rats successfully recruit LRRK2-positive monocyte-derived macrophages into the midbrain, ostensibly driven in part by responses and chemokines secreted by resident microglia, astrocytes, and other macrophages, IFNγ stimulation in isolated mouse monocyte-derived macrophages does not markedly stimulate LRRK2 expression and activity as it does in human-derived cells. These results may have implications for understanding LRRK2 function in some mouse models where monocyte-derived IFNγ stimulation drives distinct endophenotypes. Importantly, the LRRK2 kinase inhibitor MLi2 blocks α-synuclein fibril induction of LRRK2 in monocyte-derived macrophages, implicating LRRK2 kinase activity in a feedforward pathway responsive to pathogenic α-synuclein that might be interrupted with LRRK2-directed therapeutics. Future studies will be required to identify the receptors that might interact with α-synuclein fibrils in monocyte-derived macrophages, such as Lag3 (Mao et al., 2016), and the nature and consequences of LRRK2 stimulation of expression and kinase activity in these cells, and the types of signaling pathways that become activated.

## Materials and methods

### Animal and human approvals

All study protocols were approved by local Institutional Review Boards and Institutional Animal Care and Use Committees.

### Rodent strains

Strains include Cx3cr1^CreER^ (JAX Stock # 021160), ROSA26-tdTomato mice (JAX stock #007905). Cx3cr1 reporter knock-in (JAX Stock # 005582), CCR2 reporter knock-in (JAX stock #017586), R1441C-LRRK2 knock-in (JAX Stock # 009346), WT-LRRK2 mouse-BAC transgenic mice (JAX stock #012466) and C57BL/6J control mice (JAX stock #000664). Mice were genotyped according to published protocols by the depositing authors (available from the Jackson Repository), including quantitative PCR protocols used for measuring BAC copy numbers in the WT-LRRK2 mouse-BAC transgenic mice. Rats were purchased from Charles Rivers (Sprague Dawley).

### Mouse cell culture

Mouse bone marrow-derived macrophage (BMDM) cells were generated from 3-5 months-old female mice. Bone marrow cells were flushed with a 1 mL syringe containing ice-cold sterile 1x PBS and transferred through 70 µm nylon cell strainers. Cells were centrifuged at 450 xg for 10 min at 4 °C and pellets were incubated with red blood cell lysis buffer (Invitrogen) for 30 sec. DMEM media was supplemented and cell suspensions centrifuged (450 xg) for another 10 min at 4 °C. The final pellets were dissociated with pre-warmed 10% fetal bovine serum in DMEM supplemented with 20 ng mL^-1^ mouse M-CSF (PeproTech) and plated in culture dishes. When cells reach near-confluence, the monolayers were washed with DMEM two times and TrypLE reagent (Thermo Fisher Scientific) was used to detach cells. Live cells (trypan blue) were counted, and cells plated into poly-D-lysine (Sigma) coated wells for at least two days before experimentation as indicated. All experiments are curated from at least three independent experiments from at least three different mice.

### Human cell culture

Peripheral blood mononuclear cells (PBMCs) were isolated from venous blood draws. Blood draws were processed with SepMate (Fisher) tubes and Lymphoprep (Stemcell Tech) tubes according to manufacturer’s recommendations. Monocyte-derived cells were polarized and activated according to previously published protocols (Ryan et al., 2017). For M1-polarized cells (e.g., dendritic-like monocytes), cell media included RPMI-1640 Glutamax (Life-Technologies) supplemented with 10% FBS (R&D Systems/biotechne), 1% penicillin/streptomycin (Invitrogen), Fungizone (2.5 μg mL^-1^; Life Technologies), and human GM-CSF (5 ng mL^-1^). Similar, for M2-polarized cells (e.g., monocyte-derived macrophages), media included DMEM (Invitrogen) supplemented with 1x Glutamax (Life-Technologies) and 10% FBS, 1% penicillin/streptomycin (Invitrogen), Fungizone (2.5 μg mL^-1^; Life Technologies), and human M-CSF (20 ng mL^- 1^). To polarize cells towards a monocyte-derived microglia-like phenotype (MDMi, see (Ryan et al., 2017)), serum-free RPMI-1640 Glutamax (Life Technologies) was supplemented with 1% penicillin/streptomycin (Invitrogen), Fungizone (2.5 μg mL^-1^; Life Technologies), human M-CSF (10 ng mL^-1^), GM-CSF (10 ng mL^-1^), NGF-β (10 ng mL^-1^), CCL2 (100 ng mL^-1^), and IL-34 (100 ng mL^-1^). All cytokines were purchased from PeproTech. All cells were left in culture for one week before experiments. Cells were activated with LPS (100 ng mL^-1^ E. coli O55:B5, triple-purified, InvivoGen) for 48 h. Alternatively-activated cells were generated with 20 ng mL^-1^ IL-4 and IL-13 for 48 h. All experiments were curated from at least three independent experiments from cultures from two healthy male volunteers and one healthy female volunteer.

### Intracranial stereotaxic injections in rats and mice

Intracranial injections of LPS (InvivoGen) and IFNγ (PeproTech), or rAAV2/1-A53T-human-α-synuclein (Iowa Viral Vector Core, MJFF backbone), rAAV2/1-eGFP, recombinant mouse or human α-synuclein fibrils or monomer protein (generated in-house as described below and previously (Abdelmotilib et al., 2017)), or saline controls, were conducted in ∼ 8-16 week old mice or rats as described (Abdelmotilib et al., 2017; Daher et al., 2015; Daher et al., 2014). Rodents were anesthetized with vaporized isoflurane on a gas mask fitted to a digital stereotaxic frame with an integrated warming base to maintain core body temperature (Stoelting). Custom 32-gauge needles (Hamilton) with a 110° bevel were used for injections with solutions infused using an automated stereotaxic injector (Stoelting) at a flow rate of 0.25 µL per min with the bevel of the needle facing medially. All stereotaxic coordinates were empirically derived with anterior/posterior (AP) and medial/lateral (ML) measurements referenced to Bregma and dorsal/ventral (DV) measurements referenced to the skull. Bilateral mouse injections included dorsal striatum (AP +1.0 mm, ML ±1.85 mm, DV 3.0 mm) or SNpc (AP -3.1 mm, ML ± 1.5, DV -4.6 mm). For rats, SNpc coordinates were AP - 4.65 mm, ML ±2.25 mm, DV -7.45 mm. Scalp incisions were closed by monosuture. At indicated time of sacrifice, rodents were deeply anesthetized with isoflurane and transcardially perfused with cold PBS (pH 7.4) and brains removed and dissected for flow cytometry as described below or continued on with freshly prepared 4% paraformaldehyde buffered in PBS. Brains were further post-fixed for 24 h in 4% PFA/PBS for immunofluorescence analysis, floated into 30% sucrose PBS solution for up to three days, and frozen in isopentane solution (−50 °C) and stored at -80 °C for later processing to 40 μm coronal sections on a freezing sliding microtome (Leica).

### Immunohistochemistry and Immunofluorescence

Antigen retrieval was performed on all free-floating sections with a sodium citrate (pH 6.0) incubation for 30 min. Primary antibodies were incubated for 24 h followed by secondary antibodies for 24 hrs as described. For LRRK2 detection, antibody incubations were extended for both the primary and secondary antibody incubations for an additional two days. The following primary antibodies were used: CD68 (rat monoclonal FA-11, Bio-Rad), Iba1 (goat polyclonal, Abcam, and rabbit polyclonal, Wako), mouse-LRRK2 (rabbit monoclonal (c41-2), Abcam), CD163 (mouse monoclonal ED2, Bio-Rad), pS129-α-synuclein (EP1536Y, Abcam). Stereological assessments of CD163+ cells through the SNpc were accomplished with Stereologer hardware (Olympus BX61 widefield microscope) and software using the rare object counting protocol (Stereology Resource Center, Inc.). The border of the SNpc was outlined as the counting reference space for all midbrain levels at 4x magnification. Human formalin-fixed, paraffin-embedded 10 μm-thick sections of postmortem human tissues, were developed with polyclonal rabbit antibody to LRRK2 (rabbit clone HL-2; 1:250-1:500) and hematoxylin counterstain as previously described (Hakimi et al., 2011). Post-mortem brain tissues were evaluated from three idiopathic PD cases and three controls without neurodegenerative disease.

### Immunoblotting

Tissues were collected following transcardial perfusion with cold PBS. Tissue and cells were homogenized with probe-tip sonication in RIPA lysis buffer containing 50 mM Tris (pH 7.4), 150 mM NaCl, 1% Triton, and 0.1% SDS supplemented with 1x Complete protease and PhosStop inhibitor tablets (Roche). Homogenates were centrifuged at 10,000 x*g* for ten minutes at 4°C. Supernatants were analyzed using BCA assay (Pierce) to determine total protein concentration, and was then further diluted 1:1 with 2x Laemeli sample buffer supplemented with 40 mM NaF and 10% DTT. Lysates were analyzed on 4-20% gradient mini-PROTEAN TGX stain-free gels (BioRad) and transferred to Immobilon-FL PVDF membrane (Millipore), followed by immunoblotting with indicated primary and secondary antibodies. The following antibodies were used in this study: N241/34 anti-LRRK2 (Antibodies Inc), phospho-T73-Rab10 (MJF-R21, Abcam), total Rab10 antibody (MJF-R23, Abcam), loading control Hsc70 (Cell Signaling), pS129-α-synuclein (EP1536Y, Abcam), Donkey anti-mouse 680LT (LiCor), and Goat anti-rabbit HRP (Jackson Immuno). Crescendo ECL reagent (Millipore) was used to develop chemiluminescent signals and fluorescent signals were detected using the IRdye680 channel. All signals were captured digitally using a Chemidoc MP Imaging System (BioRad). Quantifications were performed using Image Lab 6.0.1 software (BioRad).

### Small molecules and administration

Small molecule LRRK2 inhibitors PFE-360 and MLi2 were synthesized in-house as previously described (Kelly et al., 2018). TLR-4 inhibitor Resatorvid was purchased from Selleckchem. All compounds were >98.5% pure as assessed by LC/MS and NMR. Compound PFE-360 was administered to rats via oral gavage in a 0.5% methylcellulose vehicle as described (Kelly et al., 2018). For induction of Cre recombinase, ∼8-10 week-old Cx3cr1^CreER^:ROSA26^tdt^ mice were treated each with four mg tamoxifen (TAM, Sigma) emulsified in 200 μL corn oil (Sigma) and injected subcutaneously at two time points 48 h apart. The first day of tamoxifen injection was considered day one in the indicated experimental time courses.

### Flow cytometry

The following anti-mouse antibodies were purchased from eBioscience, BioLegend, BD Pharm, and Life Technologies: 7-aminoactinomycin D (7-AAD) for live-dead staining; fixable viability dye (Live/Dead Fixable Dead Cell Stain Kit, Near IR); CD45 brilliant violet (BV) 650 (clone 104); CD45 BV650 (clone 30-F11); CD45 eFluor450 (clone 30-F11); CD11b eVolve 605 (clone M1/70); Ly6G allophycocyanin-conjugated (APC) (Gr-1, clone 1A8); Ly6C eFluor(eF)450 (clone HK1.4); CD11b BV605 (clone M1/70); CD11b phycoerythrin (PE) (clone M1/70); MHCII FITC (I-A/I-E); CD4 PE-Cyanine7 (clone GK1.5); CD8a APC-Cyanine7 (clone 53-6.7). All results/cytometer acquired data were analyzed using FlowJo 10.5.2 software (BD Company).

#### Blood leukocytes

Fresh mouse blood was collected as described (Lever et al., 2019). Cells from blood were centrifuged at 300 xg for 10 min at 4 °C; red blood cells (RBCs) were lysed for 5 min at room temperature using ammonium-chloride-potassium (ACK) buffer (Life Technologies). After centrifugation, the leukocyte pellet was washed with staining buffer containing phosphate buffered saline (PBS), 0.5% w/v bovine serum albumin (BSA) (Sigma) and 0.01% w/v sodium azide (Sigma). The leukocytes obtained above were stained for 30 min on ice with antibodies, after 10 min incubation with anti-mouse CD16/32 (clone 93) to block nonspecific binding of FcγRIII/II receptors. After staining and washing, data were acquired on an LSRII cytometer (BD Biosciences).

#### Whole-brain isolated leukocytes

Deeply anesthetized (isoflurane) mice were transcardially perfused with cold PBS and whole brains were removed and digested enzymatically with 1 mg mL^-1^ of Collagenase IV (Sigma) and 20 µg mL^-1^ DNAse I (Sigma) diluted in RPMI-1640 with 10% heat-inactivated fetal bovine serum (FBS), 1% L-glutamine, and 1% penicillin streptomycin (Invitrogen) as described previously (Harms et al., 2018; Qin et al., 2016). Mononuclear cells were isolated with a 30/70% Percoll gradient and blocked with Fcγ receptor antibody clone 2.4G2. Cells were then incubated with antibodies and analyzed on an Attune NxT instrument (acoustic-assisted hydrodynamic focusing cytometer; Thermo Fisher Scientific).

#### Midbrain isolated leukocytes

Deeply anesthetized (isoflurane) mice were transcardially perfused with cold PBS and whole brains were removed into a measured cold brain block (Kopf). Equally-sized 2 mm coronal sections containing midbrain were sliced, midbrains (equal sized from each mouse) were carefully dissected and minced, tissue pieces were triturated and digested in freshly made 1 μg mL^-1^ of collagenase IV (Sigma) and 20 μg mL^-1^ DNAse I (Roche) in 1x Hanks’ balanced salt solution (HBSS) (Life Technologies) for 35 min at 37°C in a water bath with gentle intermittent shaking. The digestion was halted by adding buffer (1x PBS, 2 mM EDTA, 1% w/v BSA (Sigma)) on ice. Single cell suspensions of the digested mid-brains were prepared by sequential disaggregation by pulling the homogenate through syringe needles with progressively smaller bores (18G and 20G needles) followed by passage through a 40 μm nylon filter (Fisher Scientific). After centrifugation at 650 xg for 11 min, each pellet was resuspended in RPMI-1640 (with L-glutamine) (Life Technologies) with 1% heat-inactivated FBS (Bio-Techne). The cell pellet was applied to a 30/70% discontinuous Percoll gradient by centrifuging at 500 xg at 20°C with slow acceleration and no brake in a swing bucket rotor. Cells from the interphase were collected and washed first with 1x HBSS at 500 xg for 10 min and then with 1x PBS at 500 xg for 8 min. The leukocytes obtained were stained in buffer (1x PBS, 0.5% BSA, 0.01% sodium azide (Sigma)) for 30 min on ice with antibodies, after 10 min incubation with anti-mouse CD16/32 (clone 93) to block nonspecific binding to FcγRIII/II receptors. Cells were analyzed on an Attune NxT instrument (acoustic-assisted hydrodynamic focusing cytometer; Thermo Fisher Scientific).

### Protein expression and purification of α-synuclein

Bacterial expression plasmids encoding mouse and human WT-α-synuclein in the inducible pRK172 backbone were transformed into BL21-CodonPlus (DE3) cells (Clontech). Cell pellets were lysed in 0.75M NaCl, 10 mM TrisHCl, pH 7.6, 1 mM EDTA, 1 mM PMSF and sonicated at 70% power (Fisher500 Dismembrator) for 1 min and tubes were placed in boiling water for 15 minutes. Centrifuged samples were dialyzed against 10 mM Tris (pH 7.6) with 50 mM NaCl, 1 mM EDTA, 1 mM PMSF. Suspension was passed through a HiPrep Q HP 16/10 Column, 1 x 20 mL (GE Healthcare) on an ÄKTA pure protein purification system (Cytiva, formerly GE Healthcare) with a running buffer composed of 10 mM Tris pH 7.6, 25mM NaCl, and eluted with a linear gradient application of high-salt buffer (10 mM Tris, 1M NaCl pH 7.6). Samples containing single-band profiles of α-synuclein were identified by Coomassie staining and further dialyzed and concentrated. Final concentration of protein (monomer, undiluted) was determined by BCA assay (Pierce). Purified monomer protein passed through two to three rounds of endotoxin removal (Endotoxin removal kit, GenScript) to reach a level of <0.1 EU mg^-1^, with endotoxin levels determined using a LAL chromogenic endotoxin quantification kit (GenScript). Mouse or human α-synuclein fibrils were prepared by incubation of 7 mg ml ^-1^ α-synuclein monomer of the same origin in phosphate-buffer saline for seven days at 37°C with constant agitation. Fibrils were centrifuged at 20,000 xg for 20 min at room temperature to separate high-molecular weight forms from monomer. Mouse or human α-synuclein short fibrils and contaminating higher-molecular weight species were subjected to one-hour sonication at 30% amplitude with liquid cooling maintained at 10°C (Qsonica Q700). Sonicated fibril preparations were mixed at a ratio of ∼1:10,000 w/v with monomer at 37°C with constant agitation. Seeded fibril preparations were then pelleted at 20,000 xg for 20 min at room temperature and sonicated as before to generate short rod fibrils. Preparations were measured by dynamic light scattering, with estimated molecular weight of monomers and sonicated fibrils, diluted to 0.05 mg mL^-1^ and measured on a Titan DynaPro (Wyatt Technology) at 25°C. Data were collected and analyzed using the Dyna V6.3.4 software package, with the solvent (phosphate-buffer saline) background signal subtracted from each sample. Molecular weights of particles were estimated from intensity and mass distributions. For transmission electron microscopy, 3 μL of 0.1 mg mL^-1^ samples were applied to glow-discharged 400 mesh, carbon-only, copper grids (Electron Microscopy Sciences) and negatively stained with 0.5% uranyl acetate (Polysciences). The grids were imaged in a FEI Tecnai F20 electron microscope (Eindhoven) operated at 200 kV with nominal magnification at 65,000x and a defocus range of -1.0 μm to -1.27 μm. Images were collected on a Gatan Ultrascan 4000 CCD camera.

### Microscopy and quantification

Immunofluorescence images were captured using a Leica TCS-SP5 or a Carl Zeiss 880 AiryScan confocal microscope, and brightfield images were captured on an Olympus BX61 microscope. Images were processed using Leica LASAF software, Zeiss Zen black software, with contrast and color balance adjusted in Adobe Photoshop.

### *In silico* analysis

Species differences in transcription factor binding sites at the cis-promoter of human and mouse *LRRK2* was determined using the INTERFEROME v2.0 database of IFN-inducible genes (interferome.org, (Rusinova et al., 2013)). Search conditions included: ‘All’ Interferon Types and Subtypes AND ‘hematopoietic cells’. Results for mouse (Lrrk2, ENSMUSG00000036273) and human (LRRK2, ENSG00000188906) genes were assessed under the ‘TF Analysis’ tab which included TRANSFAC matrix match of the transcription factor and its predicted binding site. GWAS variants, meta *p* values, and OR values were curated from *PDGene* and literature searches (Fava et al., 2016; Nalls et al., 2014; Zhang et al., 2009). The Genotype-Tissue Expression Portal (GTEx IGV Browser V8, gtexportal.org) was used to visualize cis-promoter region eQTLs, relevant GWAS variants related to *LRRK2* from the portal annotated ‘Whole Blood’ tissue type. The -log_10_(p-values) of eQTLs were plotted along the human reference sequence (hg38, chr12) and the positions of relevant GWAS variants were highlighted. *P* values associated with blood eQTLs were curated from the blood eQTL browser genenetwork.nl/bloodeqtlbrowser/ (Westra et al., 2013). This data portal was also used to assess relative exon expression of human *LRRK2* (ENSG00000188906) by plotting median read counts per base across 51 exons.

### Statistics

Statistical analyses were conducted using Graphpad Prism 8.0 software. Experimental groups were tested for normality and equal variance using the Kolmogorov-Smirnov test and Levene’s test, respectively. Continuous data with more than two independent groups were evaluated for significance using a one-way ANOVA, and either a Tukey’s post hoc test to compare every mean with every other mean, or a Holm-Sidak test, as appropriate, for selected group comparisons (as indicated). Two tailed, unpaired *t*-tests were used to determine significance between two groups. All tests were performed using significance level of α=0.05 with 95% confidence to determine significant group mean differences. All groups are presented with mean values and error bars representing SEM.

## Acknowledgments

We are grateful to our volunteers, patients, and their family members for participation in our research. This work has been supported by NIH/NINDS P50-NS108675 (A.B.W), NIH/NINDS R33-NS097643 (A.B.W), NIH/NINDS R01-NS064934 (A.B.W). The work has also been supported by a NSERC Undergraduate Student Research Award (to I.E.H.) and the Canada Research Chair Program (2006-2016) and a CIHR Team Grant (2014-2019) by the Government of Canada (M.G.S.).

---

**Supplemental Figure 1.**
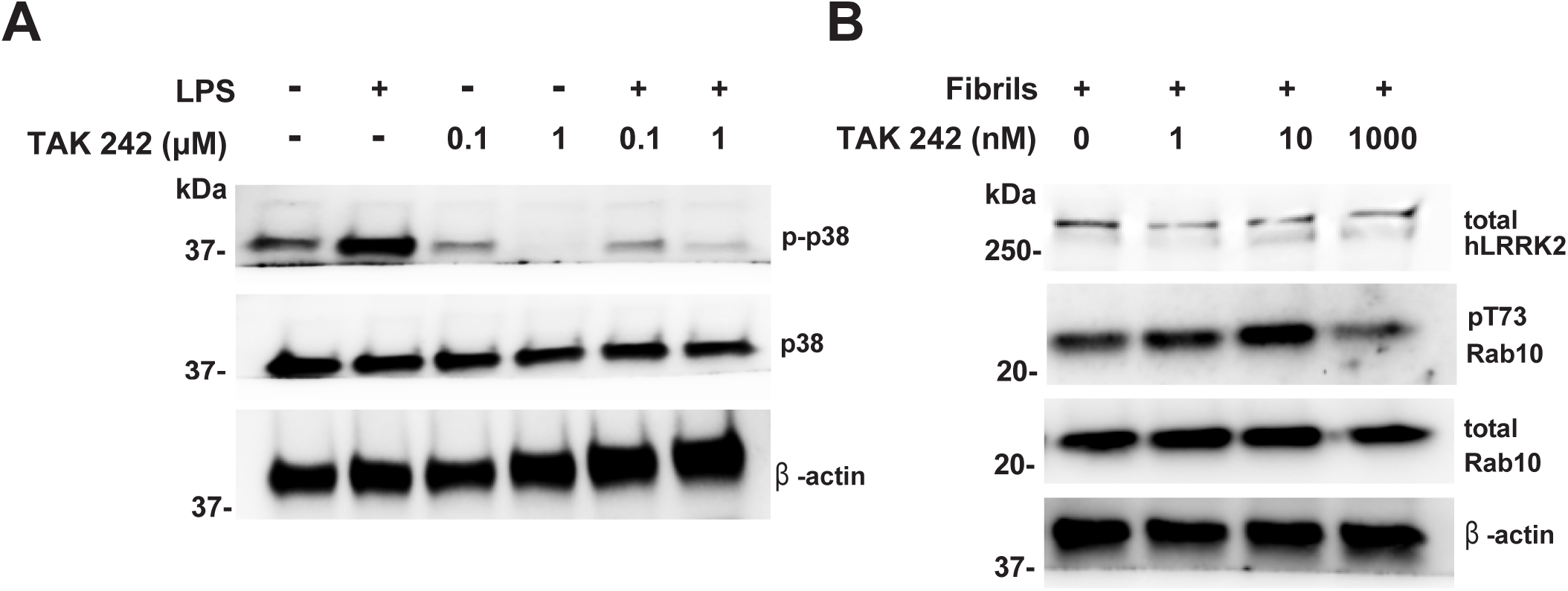
TLR4-inhibitor TAK-242 does not attenuate α-synuclein fibril induction of LRRK2 expression and phospho-Rab10 phosphorylation. **a**. Human primary monocyte-derived macrophages (M-CSF polarized) were pre-treated with TAK 242 (Resatorvid, TLR4 inhibitor, typical IC_50_ for LPS-TLR4 inhibition ∼10 nM (Ii et al., 2006)) at indicated concentration, one-hour pre-treated before LPS (100 ng mL^-1^ or 5,000 E.U. per mL) exposure. After 6-hours LPS treatment (LPS being a canonical stimulant repressed by TAK 242), cell lysates were analyzed by immunoblotting with p38 and p-p38 (downstream of TLR4 signaling pathway) to validate the efficacy of TAK-242 in our cultured cells. **b.** Human primary monocyte-derived macrophages (M-CSF polarized) were pre-treated with TAK-242 at indicated concentration one hour before the short ∼20 nm rod fibrils (1 μg mL^-1^) were administrated. After incubation with fibrils for additional 48 hours, the cell lysates were collected and analyzed by immunoblotting for LRRK2, phospho-Rab10 and total Rab10. M-CSF: macrophage colony-stimulating factor.

**Supplemental Figure 2.**
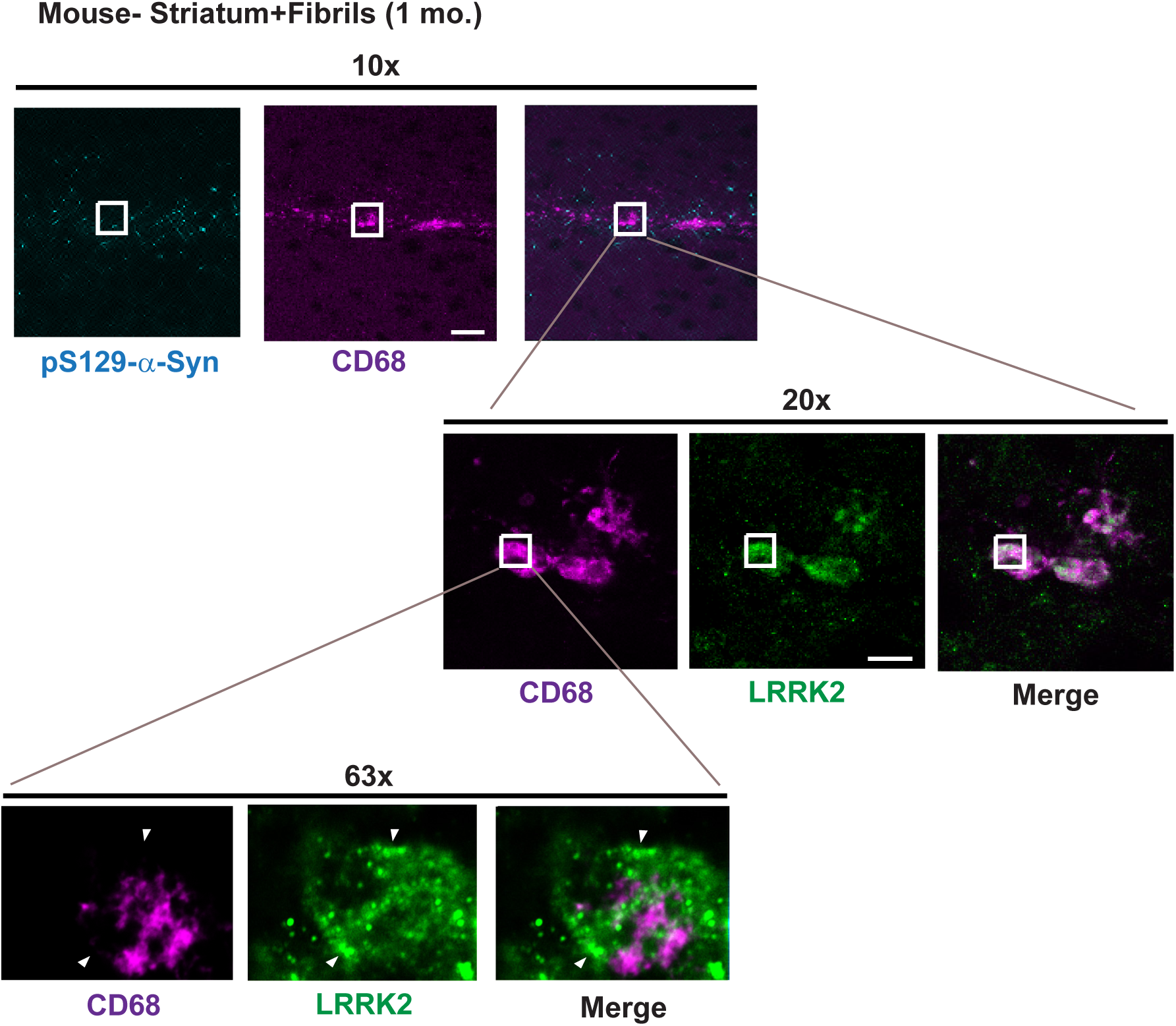
High-magnification of LRRK2 expression in CD68+ cells post α-synuclein fibril-exposure in the mouse brain. Immunofluorescence analysis of coronal sections of the mouse striatum, one month after intrastriatal injection of 10 μg of rod α-synuclein fibrils. Low magnification (10x, scale bar 200 μm) representation near the injection site in the dorsal striatum with pS129-α-synuclein staining (cyan) and a track of CD68+ cells (magenta) running through the abnormal α-synuclein. Higher magnification (20x and 63x, scale bar 10 μm) shows LRRK2 (green) with CD68 (magenta) expression, with arrowheads in AiryScan 63x images demarcating a typical endosomal-like distribution for LRRK2 inside the macrophage cytoplasm which we have noted in the past to co-localize with Rab10 endosomes closer to the plasma membrane (versus CD68). There is no significant overlap between LRRK2 and the glycoprotein CD68 inside the cell, with CD68 primarily localizing to lysosomes as well as other endosomes nearer to the Golgi.

**Supplemental Figure 3.**
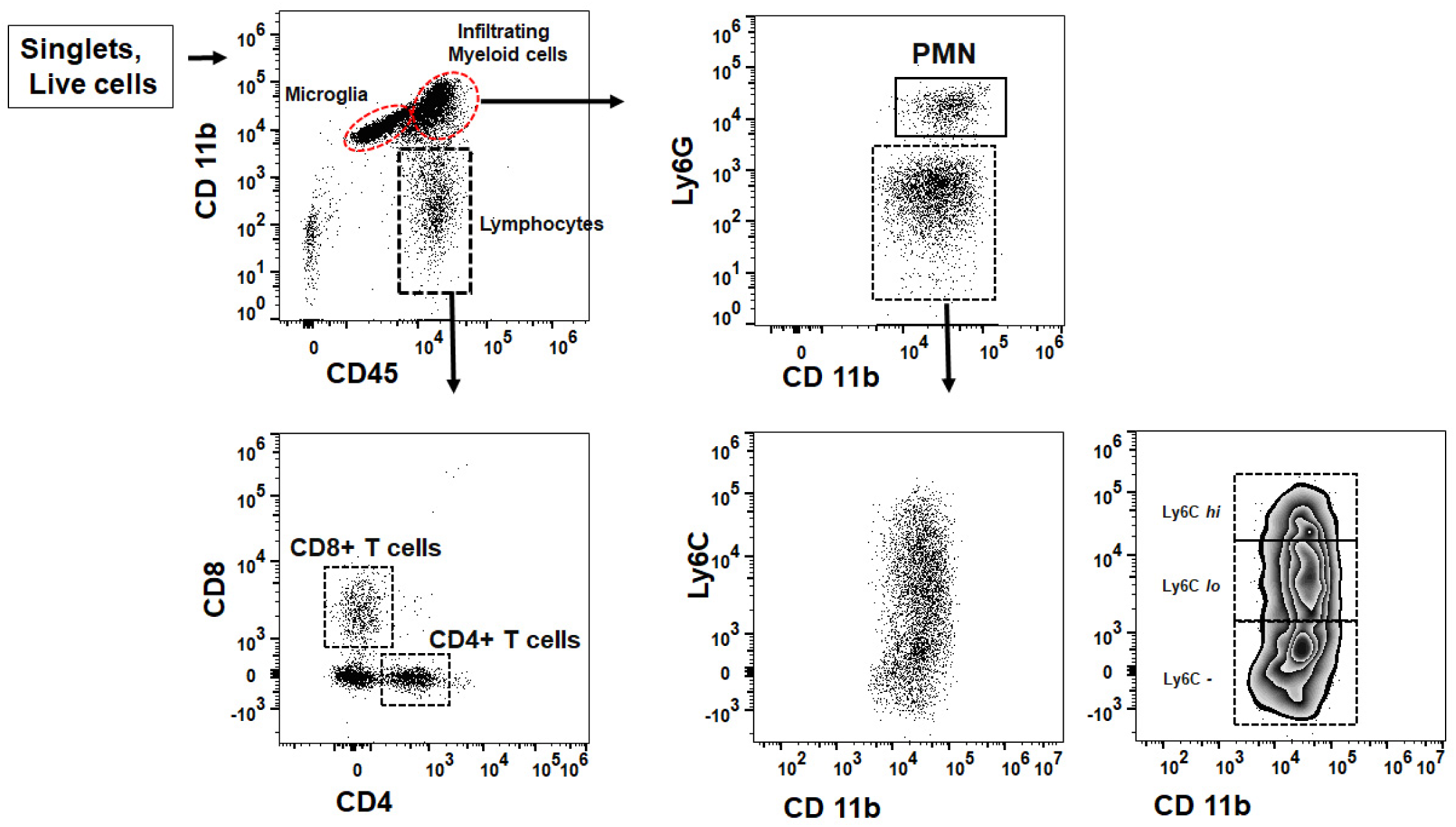
Flow cytometry gating strategy. Representative scatter plots (MFI) are shown for live CD45+/CD11b+ cells (after exclusion of debris and doublets) from mouse midbrain cell homogenates three-days after injection (bilaterally) with 10 μg of α-synuclein fibrils. Microglia are identified as CD45^int^/CD11b+ and infiltrating myeloid cells as CD45^Hi^/CD11b+. Dashed boxes represent gates, and density distribution (zebra plot) is given for Ly6C+ cells (Ly6C^Hi^ and Ly6C^lo^ cells) and Ly6C-cells. PMN: polymorphonuclear neutrophils.

**Supplemental Figure 4.**
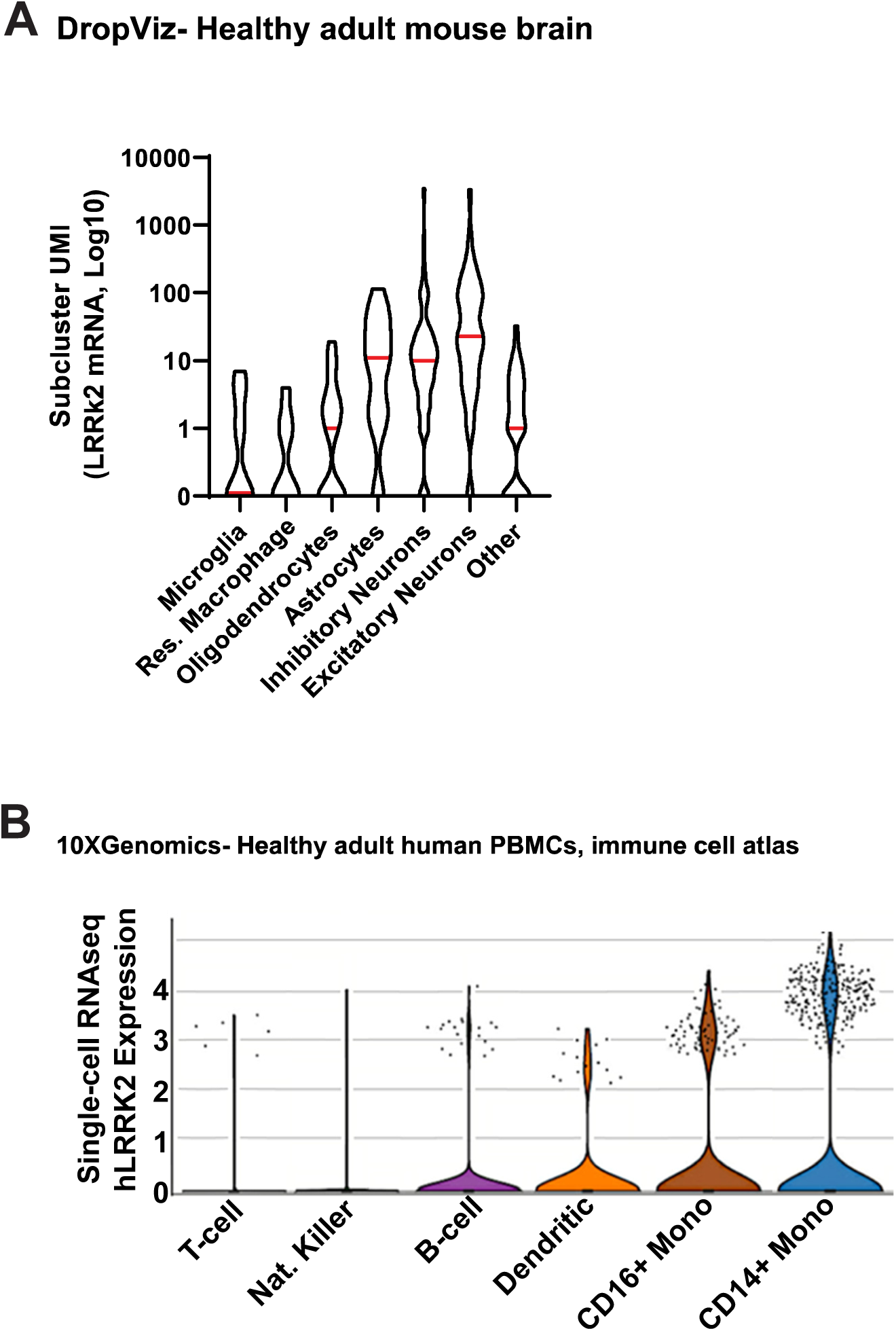
Differential LRRK2 expression in healthy brain and immune cells in the blood from healthy volunteers. **a.** Curated DropViz (DropViz.org) results for normalized mouse LRRK2 mRNA expression curated from 690,000 cells sequenced across the healthy C57BL/6 adult mouse brain. Labels represent meta-cell averages of subclusters of similar cell types through the brain that include microglia, resident macrophages (e.g., perivascular, meningeal, choroid plexus macrophages), astrocytes, oligodendrocytes, endothelial cells, inhibitory neurons, and excitatory neurons. The meta-cell average of ‘other’ includes cells that could not be assigned to these cell types. **b** Violin plots of normalized single-cell expression of human LRRK2 mRNA from sorted human peripheral-blood mononuclear cell isolations from two healthy donors (data.humancellatlas.org/). Meta-categories include T-cells, natural killer cells, B-cells, dendritic cells, and monocytes (further separated into CD16^Hi^ and CD14^Hi^ populations).

**Supplemental Figure 5.**
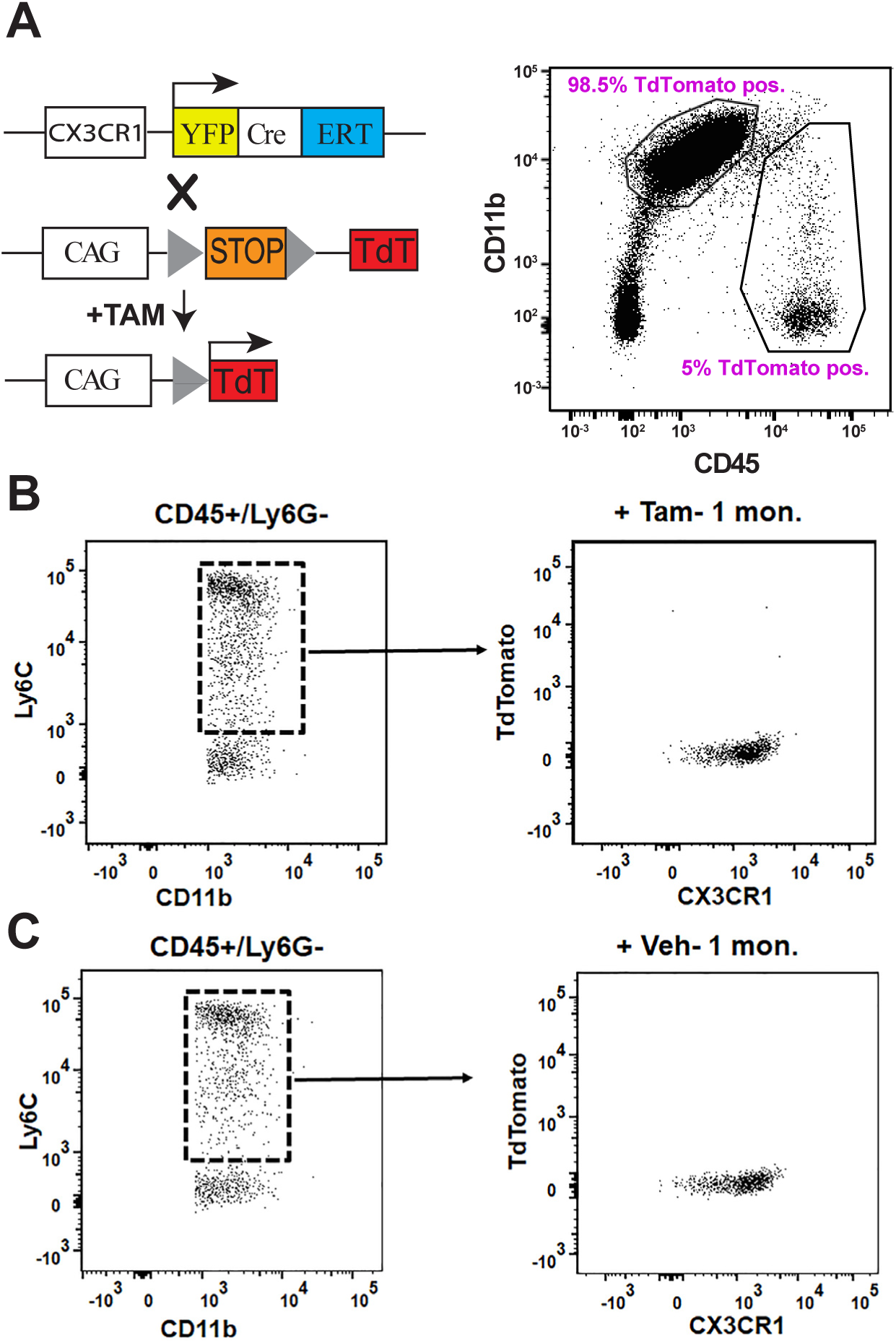
tdTomato expression in microglia and blood monocytes in double-transgenic reporter mice. **a.** Heterozygous mice with a YFP-CRE-ER fusion protein open reading frame knocked into the mouse *Cx3cr1* gene were crossed with homozygous mice with a chicken-β-actin promoter driving a flox-stop-flox tdTomato expression cassette knocked into the mouse *ROSA26* gene to generate double heterozygous mice. Transgenic mice (CX3CR1-Cre^Ert2^ / Flx-Stop-Flx-tdTomato) were injected with tamoxifen to permanently label CD45^int^/CD11b+cells (i.e., microglia) as evidenced by flow cytometry analysis (MFI) of whole-brain cell suspensions evaluated two months after tamoxifen injections. Live cells were gated by CD11b and CD45 expression, microglia were identified in whole-brain suspensions as CD45^int^/CD11b+ and the indicated populations of leukocytes (CD45^Hi^/CD11b-) were also evaluated for tdTomato expression. A representative scatter plot is shown. Mouse blood evaluated for Ly6C, CD11b, and tdTomato (MFI) in CD45+/Ly6G-cells by flow cytometry, one-month **b.** post-tamoxifen or **c**. vehicle (corn oil) injection. Representative dot plots (MFI) highlight tdTomato positive expression (from the dashed-line boxes/gates representing Ly6C+ monocytes) demarcating cells with Cx3cr1-Cre expression (n=3 mice/group; 2 males,1 female each for the vehicle and tamoxifen group).

**Supplemental Figure 6.**
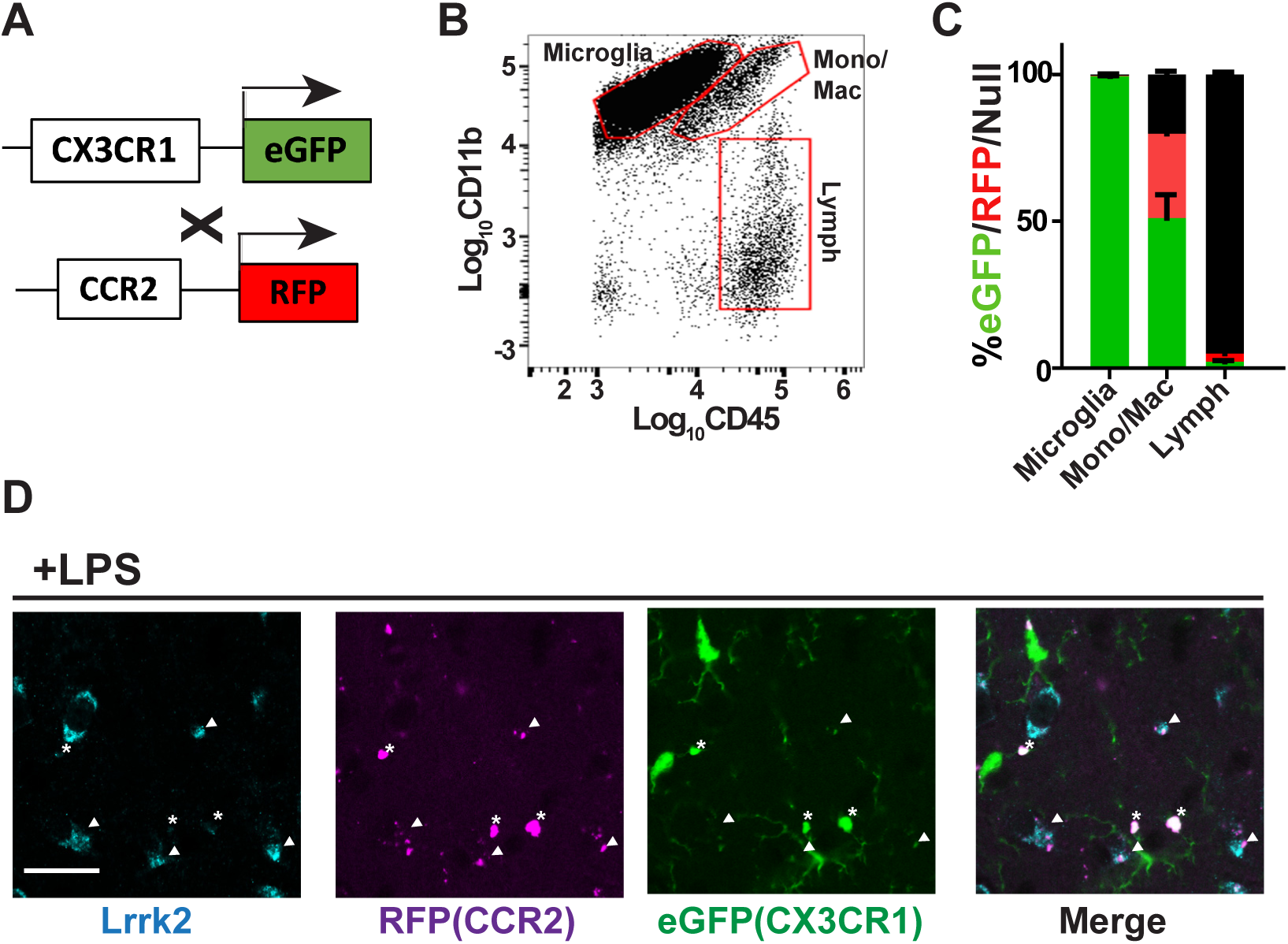
LRRK2 expression in double-transgenic monocyte/macrophage reporter mice. **a.** Double heterozygous adult mice with eGFP knock-in to the mouse *Cx3cr1* locus and RFP knock-in to the *CCR2* locus were injected into the dorsal striatum with a combination of 40 ng mouse IFNγ and LPS (10 μg, 10,000 E.U.). **b.** 48-hours after injection, cell homogenates were generated from the frontal cortex and striatum and live singlet CD11b+/CD45+ cells analyzed (MFI) for, **c.**, eGFP and RFP expression. Red boxes represent gates used to evaluate epifluorescence signal, and results are representative of three male mice analyzed that demonstrate near-complete labeling of microglia, and a mixed proportion of CD45^Hi^ cells positive for RFP (∼30%), eGFP (∼50%), or neither (∼20%). Neither eGFP nor RFP expression was detected in lymphocytic populations as expected. Column graphs show group means as a proportion of total CD45^Hi^ cells, with error bars representing S.E.M. **d.** Representative immunofluorescence images 48-hours after LPS injection in the mouse midbrain for LRRK2 (cyan), epifluorescence from RFP (magenta), and epifluorescence from eGFP (green), with merged high magnification image. Arrowheads show typical LRRK2+ macrophage-like cells that show traces (weak positive, or negative) of both RFP and eGFP, in contrast to asterisks nearby cells that are highly positive for CCR2 (but lack strong or any LRRK2 expression).

**Supplemental Figure 7.**
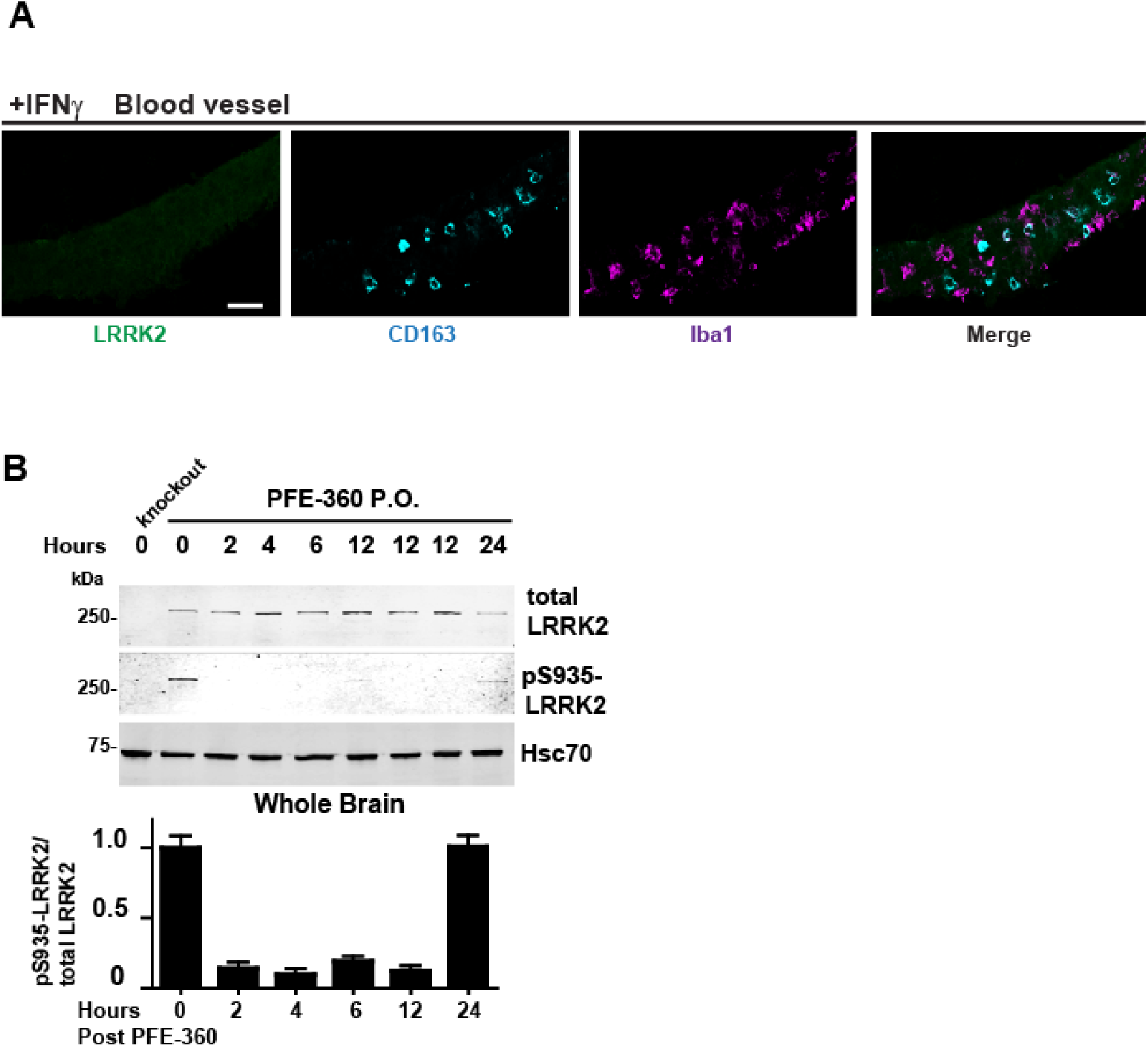
Marker analysis of blood vessel immune cells and validation of LRRK2 kinase inhibition in the rat brain with PFE-360 single-dose oral gavage. **a.** Representative immunofluorescence of CD163 (cyan) and Iba1 (magenta) in a transverse cross section of a large meningeal blood vessel, highlighting typical immune cell distribution 48-hours after injection of a combination of 40 ng mouse IFNγ and LPS (10 μg, 10,000 E.U.) into the rat midbrain. CD163+ cells were all Iba1 weak. Scale bar is 20 μm. Significant LRRK2 expression could not be resolved in these cells compared to parenchymal CD163+ cells from the same brain (Figure 4). **b.** Representative immunoblots for measures of the LRRK2 kinase inhibitor pharmacodynamic ratio (pS935-LRRK2 / total-LRRK2 ratio) as calculated by LICOR analysis, with lysates from rat frontal cortex procured two hours post oral gavage of PFE-360 (single dose, 20 mg kg^-1^). Column graph shows group means with error bars representing ± SEM.

**Supplemental Figure 8.**
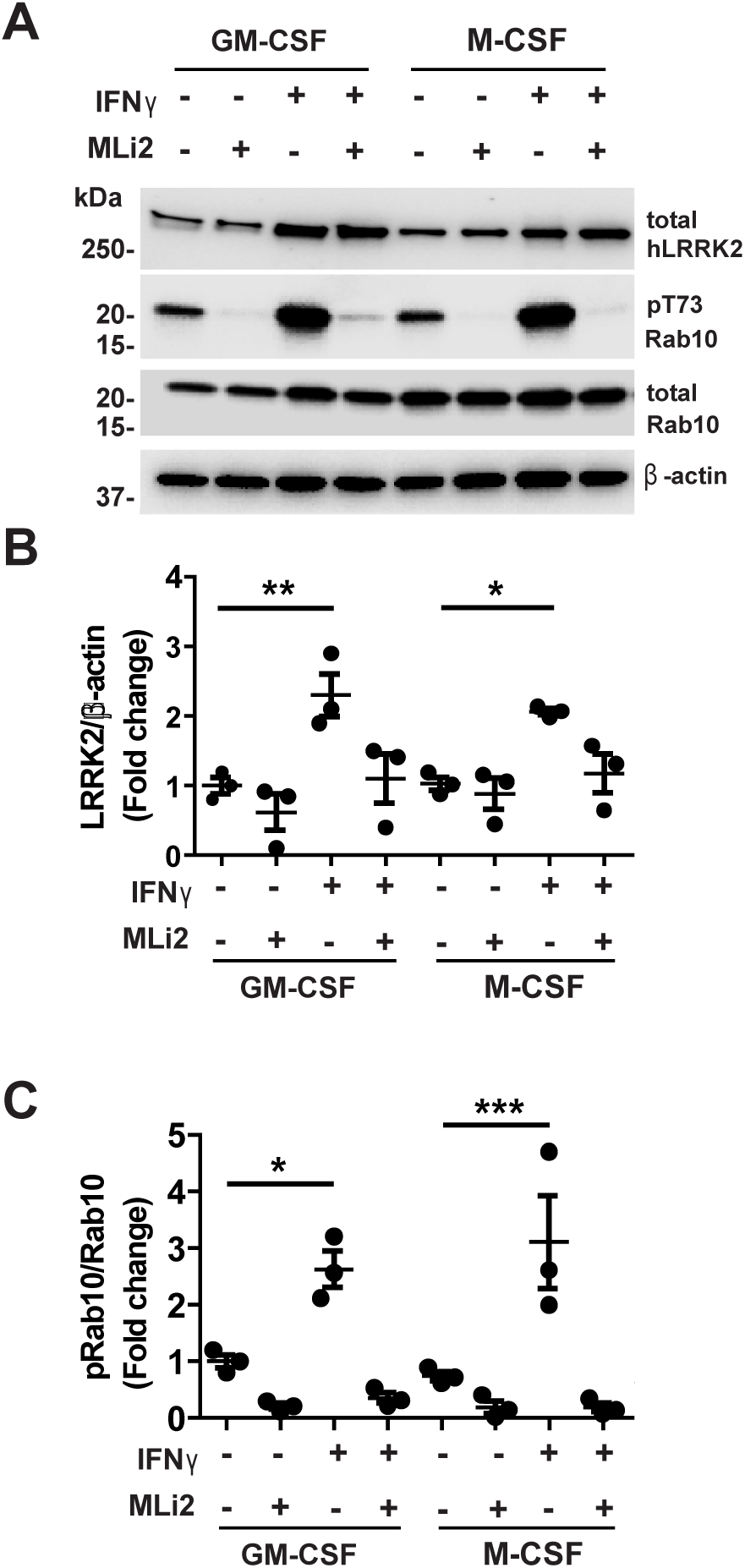
Validation of MLi2 (LRRK2 kinase) inhibition of IFNγ induction of phospho-Rab10. **a.** Peripheral blood monocytes were collected from heathy blood donors and cultured with different polarization media to induce dendritic-like cells (GM-CSF) and macrophages (M-CSF). Differentiated cells were treated with IFNγ (20 ng mL^-1^) with or without the treatment of the selective and potent LRRK2 kinase inhibitor MLi2 (cells treated at 100 nM of MLi2). Representative immunoblots show near-ablation of phospho-Rab10 levels due to MLi2 treatment, without disruption of total LRRK2 expression or total Rab10 expression, suggesting phospho-Rab10 is dependent on LRRK2 kinase activity in these cells. **b., c.**, Graphs show lines as group means with error bars representing ± SEM; significance is assessed by one-way ANOVA with Tukey’s post hoc test, *p<0.05 and ***p<0.005. M-CSF:macrophage colony-stimulating factor; GM-CSF:granulocyte-macrophage colony-stimulating factor.

